# Metastatic cell plasticity is maintained by SOX9 function during mitosis in breast cancer

**DOI:** 10.1101/2025.01.24.633933

**Authors:** Sven Beyes, Daniela Michelatti, Maria Luce Negri, Leonardo Morelli, Enrico Frigoli, Luca Fagnocchi, Alessio Zippo

## Abstract

Disseminated tumor cells (DTCs) contribute to metastasis formation by adapting to the hostile environmental conditions encountered at the seeded secondary organs. This phenotypic plasticity is supported by the epigenetic reprogramming of cis-regulatory elements (CREs), representing a hallmark of metastatic cells. DTCs can enter a reversible state of cancer dormancy, resulting from their ability to enter quiescence after mitosis and escape immune surveillance. In this respect, mitosis may represent a window of opportunity to increase DTCs’ fitness by priming CREs, ensuring the reactivation of a dormancy-related transcriptional program while preserving the epigenetic information. By profiling the chromatin changes occurring during the early steps of mitosis, we found that DTCs experience fluctuations in chromatin accessibility at CREs, which are bookmarked by pro-metastatic TFs. Among these, the pro-dormancy TF SOX9 remained associated with mitotic chromatin, facilitating transcriptional reactivation upon exiting mitosis. By dissecting the consequences of degrading SOX9 before mitosis, we found that its priming of dormancy-related CREs, combined with the recruitment of the Mediator kinase module, is instrumental for metastatic cells to enter quiescence and escape NK-mediated immune surveillance. These findings indicate that cancer dormancy is primed during mitosis by pro-metastatic TFs.

## INTRODUCTION

Metastasis is a multistep process involving cell invasion and dissemination from the primary tumor to distal sites where disseminating tumor cells (DTCs) evade the immune system, leading to overt metastasis^1, 2^. Successful progression through the metastatic cascade relies on cancer’s intrinsic and extrinsic factors, which limit the outgrowth of metastatic cells. DTCs are challenged by multiple unfavorable conditions that constrain their survival at distal sites, imposing the acquisition of phenotypic plasticity to adapt to foreign environmental cues. DTCs are exposed to immune surveillance, which is mediated by the adaptive immune response and the innate immunity mediated by natural killer (NK) cells^2–5^. Among other mechanisms, DTCs evade immune surveillance through cancer dormancy, a reversible condition in which cancer cells dynamically fluctuate from proliferative to quiescent states, reaching an equilibrium with the immune system^3–7^. Recent results indicate that tissue microenvironment and developmental signaling, which control the quiescence of somatic stem cells, also modulate cancer dormancy^8,9^. Besides these, cancer cell-intrinsic properties define the responsiveness of DTCs to these cues. This is exemplified by the epigenetic reactivation of downstream effectors such as the hormone nuclear receptor NR2F1 and the transcription factor (TF) SOX9, which are responsible for inducing cancer dormancy^5,10^. E3^11–14^. It has been shown that epigenetic reprogramming of cis-regulatory elements (CREs), including enhancers, represents a common feature that distinguishes the phenotypic plasticity of metastatic cells^12,13,15–18^. Despite these, how the epigenetic information is transmitted to the progeny during cell division while preserving phenotypic plasticity has not yet been addressed.

During mitosis, the activity of TFs is challenged by chromosome condensation, which reduces chromatin accessibility and is accompanied by loss of long-range interactions^19–21^. These chromatin rearrangements combined with the disengagement of the RNA Pol II results in a global reduction of transcription, with only a fraction of genes being actively transcribed during mitosis^22^. However, shortly after the completion of mitosis, the daughter cells reactivate the original transcriptional program, thus maintaining cell identity while preserving cell responsiveness to the environmental stimuli. To explain how epigenetic memory is transmitted it has been proposed that depositing “bookmarks” on mitotic chromatin preserves some chromatin features that facilitate transcription reactivation in post-mitotic cells. Considered as bookmarking factors are histone post-translational modifications (PTM)^23–26^, maintenance of chromatin accessibility at promoters^27,28^, TFs^19,24,29–31^, chromatin remodelers and epigenetic regulators^32,33^. Interference with any of these factors has been shown to attenuate transcriptional reactivation, affecting the maintenance of stem cell identity and cell lineage commitment^19,24,30,33^. Whether a similar mitotic bookmarking mechanism controls the plasticity of metastatic cells and their potential to enter quiescence has not been investigated. Indeed, as the choice of entering a quiescent state occurs early in G1, shortly after cell division, it suggests that metastatic cells are primed for responding to stimuli that activate a dormancy-related transcription program after completion of mitosis. In this work, we investigated whether this priming is preserved during mitosis and which bookmarking factors are involved in this process. We set out to decipher the role of metastasis-driving TFs and their differences in chromatin binding during the early phase of mitosis. We dissect the consequences of degrading the pro-dormancy TF SOX9 prior to mitosis. We found that its binding to enhancers, along with the recruitment of the Mediator kinase module (MKM)^34–36^, is instrumental for metastatic cells to enter quiescence and withstand NK-mediated immune surveillance. These findings indicate that the metastatic cell plasticity is bookmarked on mitotic chromatin by pro-metastatic TFs.

## MATERIA AND METHODS

### Cellular models

Xenograft (XD)- and Metastasis-derived cells (MD) were cultured at 37 °C and 5% CO2 in 1:1 DMEM/F-12 medium (gibco #11320-074) supplemented with 100 U/mL Penicillin/Streptomycin (gibco # 15140122), insulin (Merck, #I6634), EGF (DBA, #AF-100-15), hydrocortisone (Voden, #74144), B-27 Supplement (Gibco # 17504044) and 20 ng/ml human FGF-basic (DBA #100-18B). For G2/M arrest and release experiments, 2×10^5^ cells / ml were seeded one day prior to the addition of the inhibitor. NK-92 cells were a gift from F. Facciotti and were cultured at 37°C in RPMI (Life Technologies #21875034) supplemented with 10% FBS and with 1,000 U ml−1 IL-2 (DBA #200-02). Cells were tested for mycoplasma contamination and resulted negative.

### Derivation of stable cell-lines

Lentiviruses were generated by co-transfection of sub-confluent HEK293T cells with 9 of VSVG, 12.5 μg packaging vector pMDL, 6.25 μg pREV and 32 μg plasmid DNA of interest through CaCl transfection. HEK293T medium was changed to growth medium 12 h after transfection and lentivirus was collected 48 h later. Viral supernatants were filtered through a 0.22 mm PVDF syringe filter, concentrated by ultracentrifugation and stored at -80°C. Titration of viral particle concentration was evaluated by transduction of HEK293T cells with serial dilution of the viral preparation and assessment of the transduced cell percentage through acquisition at FACS Canto A. Viral titer was calculated as ((transduced cells N)*(( positive cell %/100)* (virusdilution)))/ ml of medium. Based on the transducing unit (TU)/mL calculated through viral titration, the viral volume used for infections was calculated based on the multiplicity of infection (MOI) as follows: TUtotal = (MOI x Cell Number)/Viral titer (TU/μL).

### Mitotic synchronization

For the G2/M border arrest, cells were incubated for 12 hours with 8 µM RO-3306 (Merck, #217699). To release cells from the arrest, they were washed three times with fresh, prewarmed medium, released into fresh medium free of inhibitors and harvested at the indicated timepoints.

For the ATAC-seq setup to capture the early timepoints during mitosis, cells were washed three times with medium containing 200 nM nocodazole (BioTrend, BN0389) and released into fresh medium containing 200 nM nocodazole and further cultured for 1 h.

### PROTAC-mediated SOX9 degradation

For the experiments using the Halo-PROTAC3 (Promega, GA3110), cells were incubated with RO-3306 for 12 hours. After 8 hours of incubation, 500 nM PROTAC were added to the medium for a total incubation of 12 hours. To release cells from the arrest and remove the PROTAC, cells were washed three times with fresh, prewarmed medium and released into fresh medium free of inhibitors and harvested at the indicated timepoints.

### NK cell-mediated cytotoxicity assay

Human activated NK-92 cells were co-cultured with *r*SOX9 tumoroids. Prior to co-culture, *r*SOX9 tumoroids were arrested at the G2/M border through incubation for 12 hours with 8 µM RO-3306 (Merck, #217699); 8h after addition of RO-3306, cells were further incubated with either 500nM Halo-PROTAC3 (Promega, GA3110) or 33nM CDK8i (SEL120-34A, MedChemExpress) for the remaining 4 hours. To release cells from the arrest and wash out the PROTAC ligand and CDK8i, they were washed three times with fresh, prewarmed medium and then seeded with NK92 cells at different effector:target ratios for 4 h at 37 °C. Following incubation, cells were stained with CellMaskTM DeepRed reagent (Life Technologies #C10046) following manufacturer’s instruction and acquired on a FACS Canto A flow cytometer and analyzed with FlowJo (BD Biosciences, v.10.5.3). Specific NK cell killing was measured for cycling and quiescent cancer cells as follows: % specificlysis=% CellMask-(dead) targets−%spontaneous CellMask-targets

### Immunofluorescence

Cells were fixed for 15 min at room temperature with 4% paraformaldehyde (Sigma-Aldrich #158127) and washed three times with PBS. Cells were trypsinized to disaggregate spheroids and trypsin was then inactivated with 10% FBS-PBS. 1 *10^5^ cells were used for each in-suspension staining. Staining was performed in 96W U-bottom plates (Corning) according to the following conditions: permeabilization and blocking with PBS/5% Goat serum (#11475055 Fisher Scientific)/1% BSA (#126579 Millipore)/0.3% Triton X-100 (blocking solution) for one hour at room temperature in constant agitation, followed by incubation with primary antibody for p27 in agitation overnight at 4°C. Cells were washed three times in PBS 1x and incubated with secondary antibodies (goat-anti-mouse or -rabbit coupled to Alexa-488 or -647 (Invitrogen), 1:500) and 1:1000 DAPI for 1 h at room temperature, in constant agitation. Cells were washed three times with 1x PBS, pelleted cells were then resuspended in 5µL mounting medium (Fluormount G # 00-4958-02 Life Technologies) and spotted on support glasses and then covered with coverslips.

### FACS analysis

To determine the cell cycle state of cells, cells were trypsinized and spun down . Cell pellet was washed once with 1x PBS, resuspended in 1 .1 ml 1x PBS and followed by 2.1 ml ice-cold 100% ethanol was slowly poured in while vortexing. For washing, 1x PBS was added twice to the fixed samples. To permeabilize cells, pellets were resuspended in 1x PBS containing 0.25% Triton X-100 and kept on ice for 15 min. Samples were spun down, supernatant was removed and pellets were resuspended in 1:200 antibody dilution (anti-phosphoHistone3-Ser10, #9701, Cell Signaling) in 1x PBS +1% BSA. Samples were incubated at RT for 90 min while gently shaking. 1 ml of 1x PBS + 1% BSA was added to the samples and then spun down. Supernatant was removed and pellet was resuspended in 80 µl 1:250 secondary antibody solution (anti-rabbit Alexa-488, LifeTechnologies) in 1x PBS + 1% BSA for 30 min at room temperature in the dark. After incubation, 1 ml of 1x PBS + 1% BSA was added and the sample was spun down. Pellets were resuspended in 50 µg/ml PI solution (Invitrogen, P3566) + 5 µg PureLink RNAse A (Invitrogen, 12091-021) in 3.8 mM sodium acetate and incubated in the dark for 30 min at 37°C.

To assess only the DNA content of cells for determining the cell cycle distribution, fixed cell pellets were resuspended in 50 µg/ml PI solution (Invitrogen, P3566) + 5 µg PureLink RNAse A (12091-021, Invitrogen) in 3.8 mM sodium acetate and incubated in the dark for 30 min at 37°C. All samples were acquired at a FACS CantoA (BD Biosciences). Data was analyzed using FlowJo 10.

### Protein lysates and western blot analysis

Total protein lysates were prepared as previously described^5^. Total protein extracts were obtained as follows. Cells were washed twice with cold PBS, harvested by scrapping in 1 ml cold PBS, and centrifuged for 5 min at 250 × *g*. Harvested cell pellets were lysed by the addition of RIPA buffer 30 min at 4 °C. Lysates were cleared by centrifugation for 10 min at 14,000 × *g* at 4 °C and supernatant was collected on ice. Protein concentration of lysates was determined using PierceTM BCA Protein Assay Kit 24 (Thermo Scientific, 574 #23227), according to the manufacturer’s instructions. The absorbance was measured at λ = 595 using Varioskan LUX Multimode Microplate Reader (Thermo ScientifiC). Values were compared to a standard curve obtained from the BSA dilution series.

For Western Blot analysis, 30 μg of protein per sample were boiled and loaded on 10% polyacrylamide gels and run in Tris-HCl-Glycine pH 8.3 running buffer + 0.1% SDS. After electrophoresis, proteins were transferred to a nitrocellulose membrane. Membranes were blocked with 5% milk in PBS-Tween (0.1%) for 1 hour at RT in agitation. Membranes were then incubated at 4°C o/n in agitation with the primary antibodies diluted in 5% milk in PBS-Tween (0.1%). Washed Membranes were and then, incubated with the corresponding secondary antibody in 5% milk in PBS-Tween (0.1%), washed again. To visualize the bands, membranes were incubated with ECL prime (RPN2236, Cytiva Life Sciences) for 1 minute in the dark and then acquired using a BioRad Chemidoc.

### DNA constructs

For the generation of the Sox9-HALO expression constructs, the linker and HALO-tag fragment were amplified from the TetO-FUW-FOXA1-HALO vector (kindly provided by Kenneth Zaret, University of Pennsylvania). Amplified linker and HALO-tag were inserted into a TetO-FUW plasmid to generate a universal entry vector for cDNA addition. The amplified SOX9 cDNA from TetO-FUW-Sox9 plasmid (addgene #41080) was then inserted into the universal plasmid using Gibson Assembly (NEB technologies) to obtain the final SOX9-HALO construct.

### Live cell Imaging

For live imaging, spheroids were disintegrated using trypsine at a 1:10 dilution, spun down and resuspended in fresh medium. 1.2 * 10^5^ cells were seeded in gelatine-coated 24 wells (ibidi) 18 hours prior to acquisition. 4 hours prior to acquisition, cells were treated with 8 µM RO-3306 to increase the number of mitotic cells at the timepoint of analysis. Cells were incubated with 200 nM Halo-substrate JaneliaFluor549 for 30 minutes and then washed three times with prewarmed medium to remove both, RO-3306 and excessive Halo-substrate. Cells were released into fresh medium w/o phenol-red and acquired with the following settings 40x objective, EM Gain 30 MHz at 16-bit, EM Gain Multiplier 250 and Conversion Gain set to 1 with 500 ms and 400 ms illumination for HALO and H2B-mCherry, respectively, using a Nikon Eclipse TI2 spinning disk microscope.

### ATAC-seq

For ATAC-seq, sample preparation according to the Omni-ATAC protocol was performed . In brief, cells were collected at the indicated time points and 50k cells were spun down for 5 min at 200 x g at room temperature, and pellets were flash-frozen in liquid nitrogen. For transposition, pellets were resuspended by pipetting up/down three times in 50 µl cold ATAC-resuspension buffer + additives (RSB; 10 mM Tris-HCl pH 7.4, 10 mM NaCl, and 3 mM MgCl_2_ in water + 0.1% Tween-20, 0.1% NP-40 & 0,01% Digitonin) and incubated for 3 min on ice. Lysis reaction was stopped by the addition of 1 ml of RSB + 0.1% Tween-20. Supernatant was removed and nuclear pellets were resuspended in 50 µl transposition mix (25 μl 2× Nextera TD buffer, 2.5 μl Nextera Tn5, 16.5 μl PBS, 0.5 μl, 1% Digitonin, 0.5 μl 10% Tween-20 and 5 μl water) and transposed for 30 minutes at 37°C and 1000 rpm. Transposition was stopped on ice and by addition of 250 μl DNA binding buffer (Zymo). Transposed samples were cleaned up using the Zymo DNA Clean&Concentrator 5 kit (#D4014). Purified DNA was mixed with 2.5 μl of each, unique dual index primers i5 andi7 (IDT technologies), 25 μl 2x NEBNext High-Fidelity 2x PCR Master Mix (NEB technologies) and water to a final volume of 50 μl. Libraries were amplified with 5 min 72°C, 30 sec 98°C, 5x(10 sec 98°C, 30 sec 63°C, 1 min 72°C). After 5 cycles of amplification, 5 μl of the sample were used for a side-PCR to define the optimal number of amplification cycles, which were then used for the remaining 45 μl of the library. Amplified libraries were purified using AMPure beads (Beckman Coulter), quantified by Qubit fluorometer (Thermo Fisher cat. #Q33226) and fragment size was assessed using the 2100 Bioanalyzer (Agilent cat. #G2939BA). Libraries were pooled at equimolar ratios and sequenced on a NovaSeq6000 (Illumina) with 75bp paired-end reads.

### Nascent RNA-seq

For nascent RNA expression analysis, cells were pulse-labeled with 0.5 mM 5-ethynyluridine for 40 min at 37°C and 5% CO_2_ prior to collection at the indicated time points. After pulse-labeling, total RNA was isolated using TRIzol (Invitrogen) and purified according to the manufacturer’s protocol. Purified RNA was measured using a Nanodrop (ThermoFisher) and further processed using the Click-iT^TM^ Nascent RNA Capture Kit (C10365, Invitrogen), as described^42^. In brief, 1 ug of 5-EU incorporated purified RNA was biotinylated, biotinylated RNA was purified by ethanol-precipitation and 500 ng biotinylated RNA was pulled down using Dynabeads™ MyOne™ Streptavidin T1 beads as described. To allow for normalization of gene expression values in mitosis, custom-made spike-in RNAs 2.5e-5 ug #1 (hg19:chr7:110258409-110258798) and 2.5-e-4 ug #2) (hg19:cr13:107098286-107098585) (expression vectors were kindly provided by Kenneth Zaret, University of Pennsylvania) were *in vitro* transcribed with biotin-UTP using the MAXIscript^TM^ SP6/T7 in vitro transcription kit (AM1322, Thermo Fisher Scientific) and added to the biotinylated RNA prior to the pulldown with Dynabeads™ MyOne™ Streptavidin T1 beads.

After washing the beads, cDNA was directly generated off the beads using the Universal Plus Total RNA-Seq with NuQuant kit (#9156, Tecan) as described by the manufacturer with the following modifications: fragmentation time was increased to 10 min and after second strand synthesis, beads were collected on a magnet and supernatant was transferred to a new tube for subsequent steps. Quality of the libraries was assessed using the 2100 Bioanalyzer (G2939BA, Agilent) and the Qubit fluorometer (Q33226, Thermo Fisher), pooled and sequenced on a NovaSeq6000 (Illumina) with 100 bp paired-end reads.

### Cut&Tag

Cut&Tag was performed using the CUTANA Cut&Tag kit (EpiCypher, 23613-1101) as described in the manufacturers protocol, with the following modifications: 100 k cells per reaction were used and mixed with 1:17 of murine NIH 3T3 cells, used as a calibrator across all samples. For amplification of libraries, each sample was mixed with 2 μl unique dual index primers, 2 μl unique dual index primers i7 (IDT technologies, stocks 10 μM) and 25 μl CUTANA High Fidelity 2x PCR Master Mix, vortexed and put in a PCR cycler with the following program: 5 min at 58°C / 5 min at 72°C / 45 sec at 98°C / 5x(15 sec at 98°C / 10 sec at 60°C) / 1 min at 72°C. For determination of additional cycles needed, 5 μl of the samples were taken, mixed with 4 μl H_2_O and 1 μl 10x SYBR and run with the following PCR program: 25x(15 sec at 98°C / 10 sec at 60°C) / 1 min at 72°C. Additional cycle number was calculated to reach ¼ of the amplification plateau. The remaining 45 μl of the samples were then run for the determined cycle number with the following program: N_cycles_ x (15 sec at 98°C / 10 sec at 60°C) / 1 min at 72°C. Amplified libraries were purified using AMPure beads (Beckman Coulter), quantified by Qubit fluorometer (Thermo Fisher cat. #Q33226) and fragment size was assessed using the 2100 Bioanalyzer (Agilent cat. #G2939BA). Libraries were pooled at equimolar ratios and sequenced on a NovaSeq6000 (Illumina) with 75bp paired-end reads.

## DATA ANALYSIS

### Immunofluorescence

Confocal imaging data analyses were performed using ImageJ software. For 2D analysis, DAPI DNA dye was used to identify the nucleus and define the region of interest. The fluorescence intensity and physical parameters were determined. The values of the fluorescence intensity were background subtracted. To quantify the nuclear mean intensity of each staining, LIF files were converted to TIFF multichannel images, and then the following macro for ImageJ was applied:

> function DAnalyze(input, output, filename, lothresh, hithresH) {
>
> open(input + filenamE);
>
> run(“Duplicate…”, “title=[TO MEASURE] duplicate”);
>
> run(“Duplicate…”, “title=C_nuc duplicate channels=1”);
>
> run(“Median…”, “radius=4”);
>
> setAutoThreshold(“Default dark”);
>
> run(“Threshold…”);
>
> setThreshold(lothresh, hithresH);
>
> setOption(“BlackBackground”, truE);
>
> run(“Convert to Mask”);
>
> run(“Fill Holes”);
>
> run(“Set Measurements…”, “area mean integrated skewness area_fraction limit display redirect=[TO MEASURE]
>
> decimal=2”);
>
> run(“Analyze Particles…”, “size=size range display clear add”);
>
> close(“Results”);
>
> selectWindow(“TO MEASURE”);
>
> roiManager(“Show None”);
>
> roiManager(“Show All”); Stack.setChannel(x);
>
> roiManager(“Measure”);
>
> saveAs(“Results”, output + filename + “.csv”);
>
> close();
>
> close();
>
> close();
>
> }
>
> input = “PathInput/”
>
> output = “PathOutput/”;
>
> lothresh = lower threshold
>
> hithresh = higher threshold;
>
> setBatchMode(truE);
>
> list = getFileList(input);
>
> for (i = 0; i < list.length; i++){
>
> DAnalyze(input, output, list[i], lothresh, hithresH);
>
> }
>
> setBatchMode(falsE);

The hitresh, lotresh, and size range parameters were determined manually for each set of images.

To assess the TF enrichment on chromatin in asynchronous and mitotic cells, chromatin, visualized by H2B-mCherry, and the cell size we defined as regions of interest (ROI) using Fiji. Quantification of chromosome enrichment was performed as described using the following equation: chromatin enrichment = log2(Intensity_Chromatin_/Intensity_WholeCell_).

For analysis of re-binding kinetics of SOX9-HALO to chromatin, images were taken every three minutes and the co-localization of SOX9-HALO and H2B-mCherry, marking chromatin, was performed using the Fiji plug-in DiANA.

### ATAC-seq

Quality of samples was assessed using fastqc and MultiQC. Reads were trimmed using trimmomatic v0.39 to remove adapter reads and low quality fragments. Paired reads were aligned to human genome hg19 using Bowtie2 with the following parameters -D 20 -R 3 -N 0 -L 20 -i S,1,0.50 –X 1000. Resulting sam files were converted to bam files using samtools v1.10, duplicates were removed using samtools markdup. Peaks were called using Genrich with the following parameters -p 0.01 -a 20 -l 0 -g 50. Peaks were annotated using <u>HOMER</u> with the command annotatePeaks.

Bigwig files were generated using the bamCoverage function of DeepTools 3.1.2^56^ for merged bam files per time point and/or treatment. Scale factors for normalization were calculated using calcNormFactors for the TMM of edgeR.

Differential accessibility analysis was undertaken using the edgeR v3.20.9 and limma v3.34.9 software packages. Normalization factors were calculated with calcNormFactors using the TMM. Differential accessibility between all timepoints and treatments was assessed using the quasi-likelihood (QL) framework of the edgeR package. P-values were adjusted for multiple testing using the Benjamini-Hochberg method. Peaks with a p-value < 0.01 and an absolute fold change > 0.5 were defined as differentially accessible regions and used for subsequent analysis.

### Transcription factor motif identification

For TF footprinting analysis, all peaks defined as distal CRE (distance greater than - 1000/+100 bp from TSS) and bigwig files of merged bam files were used as input for TOBIAS^38^. As a source of TF motifs, a curated list derived from the NGS-analysis pipeline IMAGE^57^ was used. For TF footprint analysis of peaks changing upon SOX9 depletion, merged bigwig files per time point and treatment and a peak file composed of all differential distal peaks were used as input.

As an independent approach, differential distal peaks either gaining or losing accessibility upon PROTAC-mediated deletion of SOX9 were used as input for motif identification using HOMER with the following command findMotifsGenome.pl peaksA hg19 -bg peaksB -size given -p max, using one file as input and the other as background and vice versa.

### Nascent RNA-seq

Raw reads were quality controlled using FastQC and trimmed using Trimmomatic v0.31. Reads were aligned to the human genome hg19 using BowTie2) with the following parameters -D 20 -R 3 -N 0 -L 20 -i S,1,0.50 -X 1000 and resulting SAM files were further processed and duplicates were removed using samtools v1.10. ‘MultiBamSummary’ of Deeptools v3.1.2^56^was used to retrieve the counts per sample over the RefSeq-transcript model hg19 in bed format. Counts transformed to FPKM values and spike-in normalized as described^42^. Genes were filtered for an expression level in asynchronous state of FPKM > 1. For the analysis of the expression of transcription factors only, the remaining genes were merged with a table containing all true TFs derived from the human transcription factor database^58^.

GO terms analysis was conducted using the R package enrichR and the following collections of terms “GO_Molecular_Function_2021”, “GO_Cellular_Component_2021”, “GO_Biological_Process_2021”, “KEGG_2021_Human” and “MSigDB_Hallmark_2020”). Bigwig files per timepoint were generated using the bamCoverage function of DeepTools 3.1.2. and the spike-in normalized reads as source for the normalization factors.

### Peak calling Cut&Run & downstream analysis

For defining active and poised enhancers and H3K4me3 broad domains, published Cut&Run data for. Cut&Run samples normalized on NIH3T3 spike-in for SOX9, H3K27ac, H3K4me1, H3K4me3, MED1 and an IgG control was used as previously described^5^. Peaks were called using HOMER with the following parameters -style histone Cut&Run_sample –C0 –P 0.00001 –r –size 1000 – minDist 2500 –i IgG. Called peaks for H3K27ac and H3K4me1 were merged with distal peaks called for ATAC-seq to identify enhancer regions using the Genomic Ranges package in R .

### Cut&Tag

Quality of samples was assessed using fastqc and MultiQC. Reads were trimmed using trimmomatic v0.39 to remove adapter reads and low-quality fragments. Paired reads were aligned to human genome hg19 and mouse mm10 genome (spike-in) using Bowtie2 with the following parameters -D 20 -R 3 -N 0 -L 20 -i S,1,0.50 –X 1000. Resulting sam files were converted to bam files, reads on chrM and low-quality reads (-q 30) were removed using samtools v1.10. Bigwig files per timepoint were generated using the bamCoverage function of DeepTools 3.1.2. Samples were normalized for the NIH 3T3 spike-in and further normalization was carried out by using the reads of Cut&Tag samples falling into accessible regions defined by ATAC-seq.

### Publicly available data analysis

Comparison of TF clusters showing different reactivation kinetics to publicly available data was done by downloading the TCGA BRCA gene expression dataset via XenaBrowser and plot the cluster-wise expression per BC subtype. To identify potential co-factors involved in SOX9-mediated regulation of distal CRE prior to and during mitosis, BioGrid was queried for potential SOX9 interactors. To further narrow down the potential interactors, publicly available cell-cycle resolved proteomics data was filtered for potential interactors^49^. Data was visualized as described in the original publication by calculating the z-score over log2-transformed normalized values.

### Data visualization

Plots were generated using the ggplot2 (v 3.3.3) package in R. For heatmaps, the ComplexHeatmap (v 2.6.2) package in R was used Venn diagrams and UpSetR plots were generated using the R packages VennDiagram (v 1.7.1) and UpSetR (v 1.4.0), respectively. Nascent RNA-seq coverage plots were plotted using the R package Sushi (v 1.28.0). Heatmaps and average profiles of peak signals were plotted using Deeptools 3.1.2 with the commands plotHeatmap and plotProfile, respectively. Principal component analysis was done using the computeMatrix and plotPCA commands from DeepTools 3.1.2 by either using all ATAC-seq peaks or only using the differential accessible peaks identified by ATAC-seq as region files.

### Statistical Methods

Biological replicate sample sizes (n = x) and other experiment-specific details are indicated in the Figure legends. Data were plotted and analyzed using either GraphPad or R. Data are represented as the mean of multiple replicates ± SD or SEM. In all analysis, exact p-values are indicated in the corresponding Figures.

## RESULTS

### Dynamic changes in chromatin accessibility and TF binding pattern characterize prometaphase of cancer cells

We investigate whether mitosis represents an opportunity for cancer cells to increase their fitness during tumor progression and metastasis formation by augmenting their plasticity upon cell division. To address this question, we compared the dynamic changes in chromatin accessibility during mitosis among xenograft (XD)- and metastasis (MD)-derived tumoroids^5^. To capture the initial steps of mitotic structural rearrangements that possibly limit chromatin accessibility and TFs binding^19,20^, we employed a double-inhibitor block synchronizing the cells at the G2/M border with the Cdk1 inhibitor RO-3306, followed by nocodazole treatment to arrest them at prometaphase (Figure 1A). Staining for the prometaphase-specific histone mark phosphorylation of Serine 10 of Histone (H3Ser10ph) showed the successful arrest and the subsequent release of cells into both early and late prometaphase (Supplementary Figure 1A). To assess the changes in chromatin accessibility during the early time points of mitosis, we performed a calibrated Assay for Transposase-Accessible Chromatin using sequencing (ATAC-seq) on both XD- and MD tumoroids. The genomic distribution of the chromatin accessible regions in prometaphase showed enrichment for promoters and distal CREs, thus recapitulating the pattern detected in interphase (Supplementary Figure 1B). We found that in early prometaphase, chromatin accessibility decreased both at distal and proximal CREs, independently from the cancer cell types. (Figure 1A, C, and Supplementary Figure 1C). In late prometaphase, chromatin accessibility was partially restored, particularly at the level of the promoters. This dynamic chromatin reshaping at the CREs phenocopied the global changes in chromatin compaction and organization that have been previously described to occur during prometaphase^20, 37^. We then asked whether the observed changes in chromatin accessibility during prometaphase were associated with the putative binding of TFs at CREs, which was assessed by analyzing TF footprints^38^. Focusing on distal CREs, we found that upon entry in mitosis, footprint scores of CTCF, KLF family members, and BACH2 were decreased (Figure 1D, Supplementary Figure 1E, and Supplementary Table 1), in line with previous reports on mitotic bookmarking properties of TFs^29^. Interestingly, binding of TFs belonging to the SOX family such as SOX9 or CDX and HOX family members, were predicted to be enriched in early prometaphase when chromatin was the least accessible (Figure 1D, e and Supplementary Figure 1D, E), indicating a potential role of these TF families during chromatin binding in early mitosis. For example, SOX9 motifs were enriched in the early prometaphase compared to the asynchronous sample and not compared to the late prometaphase. Indeed, upon mitotic progression, we detected an enrichment for the footprint score related to TFs belonging to the MAF and AP-1 families in both MD and XD tumoroids (Figure 1E, Supplementary Figure 1E, and Supplementary Table 2). By analyzing the enrichment of TF motifs at the TSS, we identified the nuclear transcription factor Y family (NFY) as the top hit in the asynchronous and late prometaphase condition in both cell types, which confirms the role of NFY TF in bookmarking TSS and regulating mitotic gene expression^39, 19^. Of note, we did not retrieve DNA binding motifs enriched at the TSS in the early prometaphase condition (Supplementary Figure 1F, G, and Supplementary Table 2). Indeed, compared to the distal CREs, few TFs motifs were identified as being enriched at the TSS, suggesting the involvement of rather a common mechanism to maintain a certain level of chromatin accessibility and bookmarking^40,41^. Taken together, these dynamic changes in chromatin accessibility during early mitosis and the associated enrichment for TF motifs, specifically at the distal CREs, suggest a key role of TFs in bookmarking these regulatory elements.

**Figure 1.**
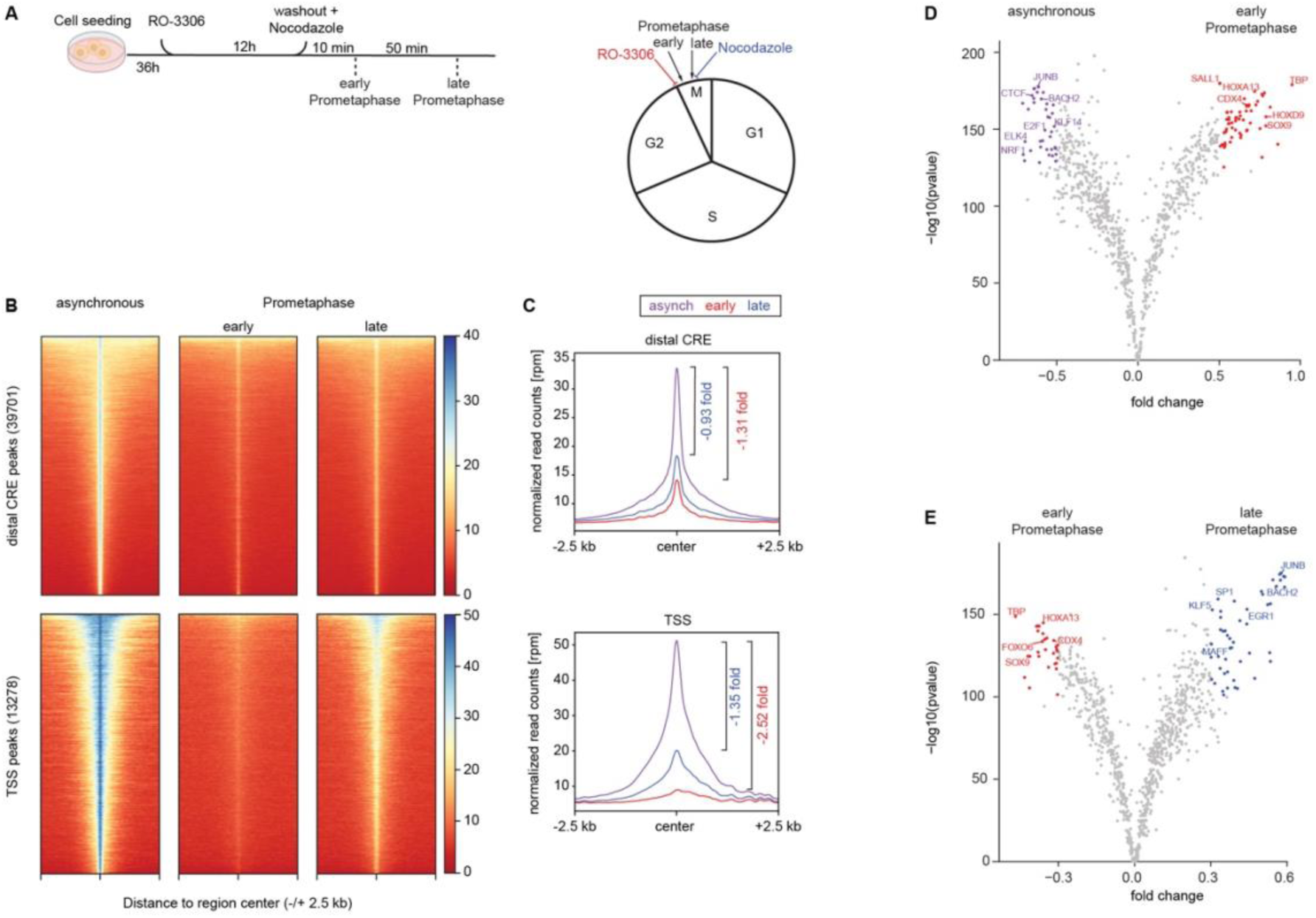
Chromatin accessibility is decreased during early mitosis. (A) Scheme showing the synchronization protocol to investigate chromatin accessibility changes during early mitosis. Cells were synchronized 12 hours with the CDK1 inhibitor RO-3306 to arrest them at the G2/M border. CKD1 inhibitor was washed out in the presence of nocodazole. Cells were harvested after 10 and 60 minutes in presence of nocodazole to enrich for cells in early and late prometaphase, respectively. (B) Heatmaps showing chromatin accessibility determined by ATAC-seq at distal *cis*-regulatory element (CRE) peaks (top) and TSS peaks (bottom), on MD tumoroids in the asynchronous condition, during early, and late prometaphase. Signal is centered on the peak center with a window of -/- 2.5 kb. Shown is the intensity of two merged replicates. (C) Chromatin accessibility profile determined by ATAC-seq at distal CRE and TSS on MD tumoroids at the asynchronous condition, during early and late prometaphase. Difference in accessibility between the asynchronous state and early and late prometaphase is depicted as log2fold change (log2FC). (D) Volcano plot showing results of TF footprint analysis of distal CRE comparing asynchronous vs. early prometaphase in MD tumoroids. Fold change is shown on x-axis, - log10(pvalue) is shown on y-axis. Significant TF motifs for both timepoints are colored and selected TFs are labeled. For full list of TFs, see Supplementary Table 2. (E) Volcano plot showing results of TF footprint analysis of distal CRE comparing early vs. late prometaphase in MD tumoroids. Fold change is shown on x-axis, -log10(pvalue) is shown on y-axis. Significant TF motifs for both timepoints are colored and selected TFs are labeled. For full list of TFs, see Supplementary Table 2.

### Expression of cancer-associated TFs is preserved during mitosis

We investigated whether the retained chromatin accessibility at distal CREs resulted from the preserved expression and activity of the specifying TFs throughout mitosis. Therefore, we measured the relative abundance of nascent transcripts by EU-RNA-seq in MD tumoroids arrested at the G2/M border or during transition through meta/anaphase (40 min.), telophase/cytokinesis (120 min.), and post-mitosis (240 min.) (Figure 2A, B). We found that entry into mitosis was associated with a global quenching of transcription that was resumed upon exiting mitosis and reached complete reactivation in the daughter cells (Figure 2C, and Supplementary Table 3)^42^. K-means clustering of nascent transcripts revealed different patterns of gene expression reactivation during mitosis, indicating functional diversity among the gene clusters (Supplementary Figure 2A, B, and Supplementary Table 3). Genes showing a spike in transcription at the G2/M border were functionally enriched for apoptosis, amino acid production, p53, and mTORC pathway (Supplementary Figure 2B, C, and Supplementary Table 3)^43^. Early reactivated genes showed enrichment for functional terms associated with protein modification, transcription initiation, and protein ubiquitination (Supplementary Figure 2D and Supplementary Table 3). Most genes reactivated upon mitotic exit were functionally enriched for biological processes linked to RNA processing, organelle formation, and nuclear lumen. In contrast, genes reactivated post-mitosis were enriched for processes linked to DNA replication and mitosis (Supplementary Figure 2E, F , and Supplementary Table 3)^42,44^. Using LMNA as a reference gene for monitoring transcription dynamics, we observed its reactivation starting from 40 min post-release, a time point overlapping with meta/anaphase, indicating a timely precise upregulation of LMNA expression to re-establish the nuclear envelope in the daughter nuclei in time (Supplementary Figure 2G)^45^.

**Figure 2.**
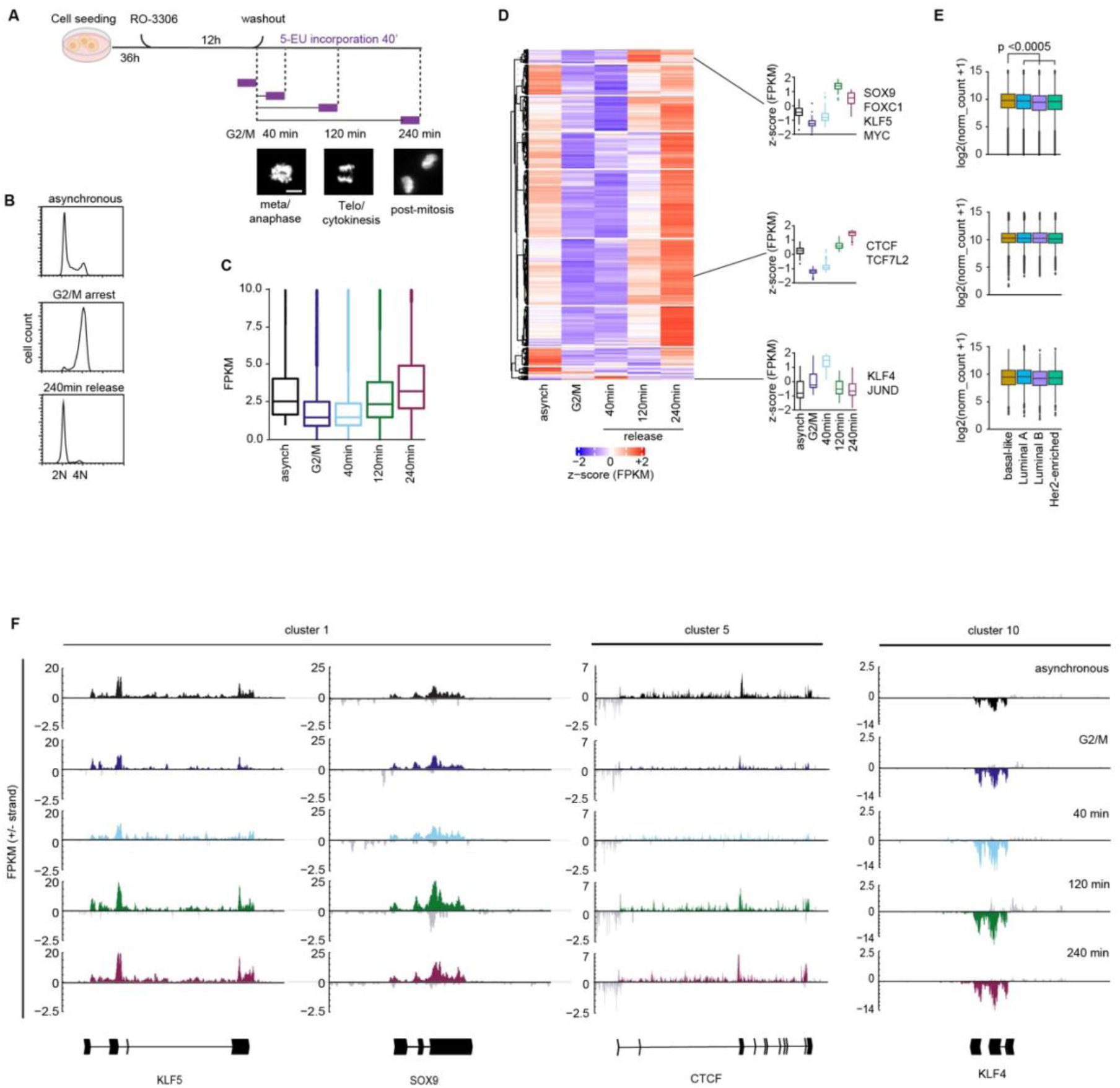
Transcription of a subset of TF is maintained during mitosis. (A) Scheme showing the experimental setup for nascent RNA-seq analysis during mitosis. Cells were synchronized for 12 hours with the CDK1 inhibitor RO-3306 to arrest them at the G2/M border. Inhibitor was washed out and cells were incorporated for 40 min with 5-Ethynyluridine (5-EU) to label nascent RNA at the indicated timepoints. Timepoints of 5-EU incorporation are labeled in purple. (B) Cell cycle distribution determined by FACS showing the arrest and washout efficiency using RO-3306 for a G2/M arrest. Timepoints shown are asynchronous, G2/M and 240min post-release from the CDK1 inhibitor. Cells were stained for DNA content using propidium iodide (PI) for determination of DNA content, as shown on x-axis. Y-axis shows cell count. (C) Boxplot showing the changes in overall FPKM of expressed genes (defined by FPKM in asynchronous cells > 1) at different time points (G2/M arrest, 40min, 120min and 240min release). (D) Heatmap showing k-means clustering in 10 clusters of TF gene expression changes during mitosis after a G2/M arrest using RO-3306, as indicated in Figure 2. Shown is the z-score of the FPKM values over the indicated timepoints per TF gene. For Genes per cluster and order of clusters, see Supplementary Table 4. (E) Boxplots showing the expression values of the genes per cluster and breast cancer subtype derived from TCGA. P-values determined by two-tailed unpaired t-test. (F) Browser view of example genes for reactivation of expression corresponding to the clusters indicated in panel D. Shown are the forward and reverse strand per timepoint asynchronous, G2/M border, 40min, 120min and 240min release. Coordinates (hg19) shown are: KLF5 chr13: 73630000-73655000; SOX9 chr17: 70112913-70128505; CTCF chr16:67586725-67678364; KLF4 chr9:110233169-110265994.

By focusing on the expression pattern of TFs, we found that most genes showed a marked downregulation upon entry into mitosis, followed by transcriptional reactivation with variable kinetics (Figure 2D and Supplementary Table 4). Indeed, we identified a cluster of cancer-related TFs whose expression was marginally reduced at the G2/M border and then increased from meta/anaphase on, reaching the highest expression level before mitotic exit (Figure 2D). This cluster was composed of TFs important for basal breast cancer identity, such as SOX9, FOXC1, and KLF5 (Figure 2E)^4,5,10^. We identified a second cluster of TFs through the same analysis, whose expression was robustly diminished during mitosis and reactivated upon mitotic exit. This gene expression pattern was represented by the Wnt-pathway effector TCF7L2 and the insulator CTCF (Figure 2D, 2F). Of note, we found that the expression of a small subset TFs, such as KLF4, was preserved throughout mitosis, suggesting a mitotic-related gene regulation pattern (Figure 2D, 2F). Considering that the expression level is not always predictive of the corresponding protein abundance, we monitored the protein levels of a subset of TFs throughout mitosis. We found that while the CTCF and MYC recovered slowly after release from the G2/M border, SOX9 protein abundance increased rapidly showing a full recovery before mitotic exit (Supplementary Figure2H, I). These results indicate that although transcription is globally shut down upon entry into mitosis, the expression of a subset of cancer-related TF is partially preserved during mitosis and further increased before mitotic exit.

### The pro-metastatic TF SOX9 is retained on mitotic chromatin

Considering the gene expression pattern of cancer-related TFs, we investigated whether they may contribute to mitotic bookmarking to facilitate transcriptional reactivation upon exiting mitosis. Among these, we focused on SOX9, which has been shown to contribute to cancer progression and metastasis formation by sustaining cancer dormancy to escape NK immune surveillance^4,5,7,10^. RNA fluorescent *in situ* hybridization (smFISH) labelling of the nascent transcript confirmed that SOX9 expression occurred during mitosis (Figure 3A). Of importance, SOX9 protein was similarly preserved, reaching the same level detected in interphase shortly after the G2/M border transition (Supplementary Figure 2H, I). Considering that the entry into quiescence occurs mainly shortly after cell division^46^, we investigated whether SOX9 contributes to this decisional step by bookmarking chromatin domains during mitosis. We employed a degron system to temporally control the protein level of SOX9 in metastatic tumoroids upon entering mitosis. Specifically, we combined the knock-down of the endogenous SOX9 transcript with the constitutive expression of a HALO-SOX9 construct that re-established its protein level in metastatic tumoroids (thereafter named *r*SOX9) (Figure 3B, Supplementary Figure3A, B). Functional characterization of *r*SOX9 tumoroids showed comparable proliferating capabilities and a similar frequency of quiescent cells (Supplementary Figure 3C-F), with respect to the native metastatic tumoroids. Furthermore, by comparing the corresponding RNA-seq profiles, we did not observe relevant changes in gene expression (Supplementary Figure 3G), indicating that the designed approach preserved the cellular functionality. Thereafter, we employed a HALO-based labeling strategy to detect the kinetics of SOX9 chromatin binding during mitosis in living cells^28^. Although the chromatin-bound fraction of SOX9 was diminished in mitosis compared to cells in interphase, it was not entirely excluded from mitotic chromatin, showing a uniform distribution within the cells (Figure 3C). To rule out whether an increased protein turnover determined the change in SOX9 distribution throughout mitosis, we measured its lifetime by live imaging and observed only a modest decrease after anaphase (Supplementary Figure 3H). By analyzing the kinetics of SOX9 redistribution during mitosis, we measured a 2-fold decrease of its association during ana/telophase (Figure 3D-F). However, shortly after cytokinesis, SOX9 rebound chromatin even before chromatin decompaction, as shown by monitoring the pattern of H2B relaxation with respect to interphasic chromatin (Figure 3D-F). These results point towards a role of SOX9 in mitotic bookmarking, as it remained partially bound to mitotic chromatin and rapidly re-associated before chromatin decompaction.

**Figure 3.**
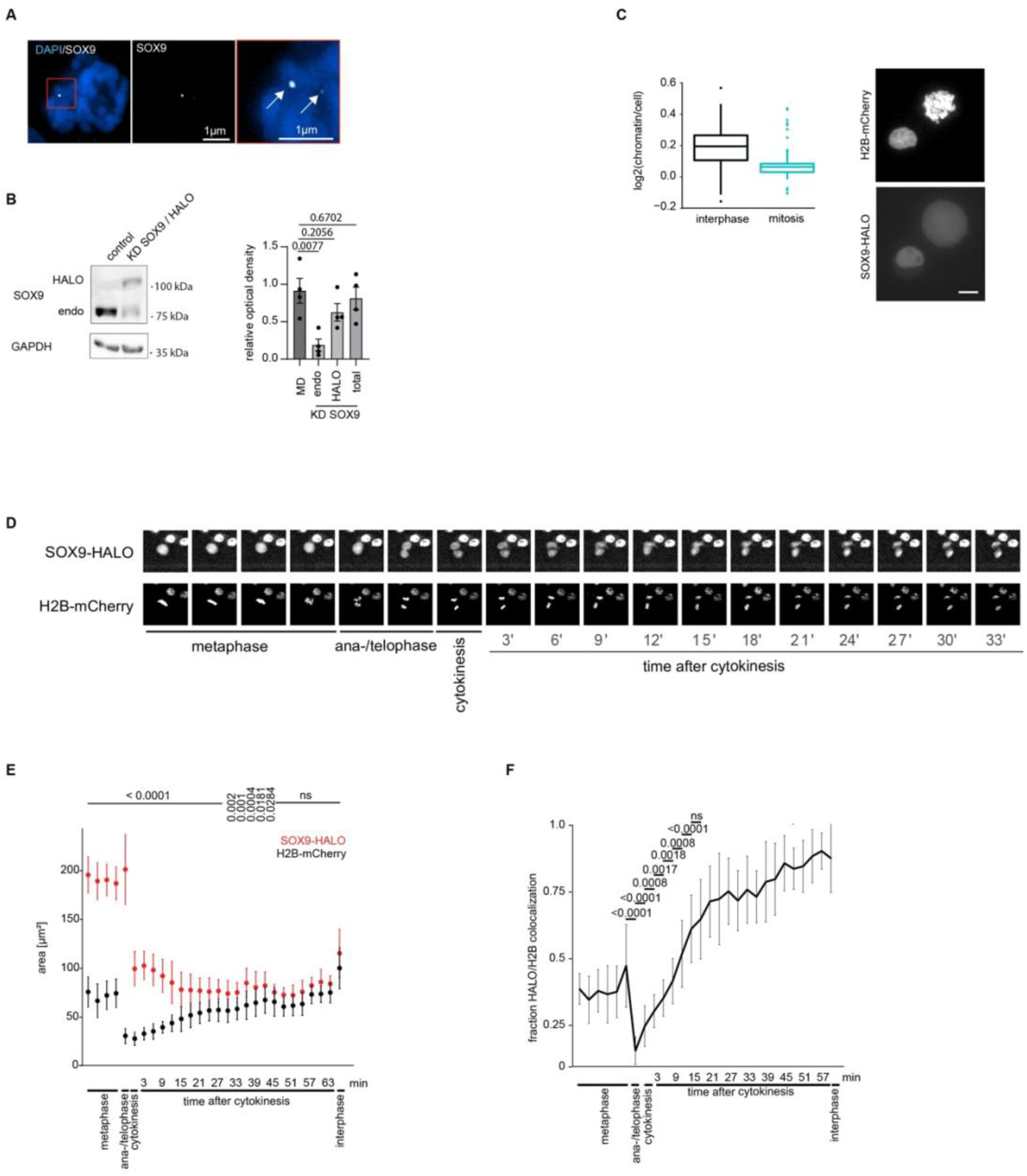
SOX9 is expressed and not degraded during mitosis. (A) Representative images of smRNA-FISH for *SOX9* on mitotic cells depicted among asynchronous metastatic tumoroids. Nuclei are stained with DAPI. Arrows indicate loci of active transcription. Scale bar: 1 µm. (B) Immunoblot for SOX9 in parental and rSOX9 tumoroids (labeled as KD SOX9 / HALO) cells. The endogenous SOX9 and SOX9-HALO were detected using a SOX9-specific antibody. GAPDH was used as loading control. Right panel: Quantification of SOX9 protein levels in the two cellular models, depicted in three independent biological samples. P-values are reported in the Figure. (C) Representative images and quantification of the chromatin-associated fraction of SOX9-HALO in asynchronous (116) and mitotic (53) cells. Left: Chromatin labeled by H2B-mCherry signal, HALO staining for SOX9-HALO. Scale bar. 10 µm (D) Representative images of SOX9 localization and distribution during mitosis. Images were taken every three minutes after release from RO-3306-mediated G2/M arrest over a time course of 1 hour. (E) Quantification of the area of SOX9-HALO and H2B-mCherry signal area during mitosis, indicating the different phases of mitosis based on changes in nuclear size determined by H2B-mCherry. Images were acquired from three independent biological samples. P-value is shown for each timepoint, comparing HALO- and H2B-mCherry-signal area. P-values determined using two-tailed t-test are reported in the Figure. (F) Quantification of the chromatin-associated fraction of SOX9-HALO during mitosis, as depicted by live imaging. Images were taken from three independent biological samples every three minutes and different mitotic phases are indicated. P-values determined using two-tailed t-test are reported in the Figure.

### SOX9 maintains chromatin accessibility during mitosis

To analyze the contribution of SOX9 to mitotic bookmarking, we employed a HALO-tag-based protein degradation strategy to temporally control its protein abundance throughout mitosis^47^. Shortly incubating the *r*SOX9 tumoroids with the HALO-PROTAC3 allowed us to degrade the exogenous HALO-SOX9 protein and weaken the SOX9 feed-forward regulatory loop, which resulted in the downregulation of the remaining endogenous SOX9^48^ (Figure 4A, B and Supplementary Figure 3I, J). By adopting this strategy, we depleted SOX9 protein four hours before releasing the cells from the G2/M blockage and measured its contribution to the maintenance of chromatin accessibility at the distal CREs by ATAC-seq (Figure 4C). Differential analyses highlighted that SOX9 perturbation affected mostly distal regulatory elements, while we did not observe relevant changes in chromatin accessibility at promoters, in line with SOX9 preferential binding at enhancers^5^. By performing cluster analyses, we identified distal CREs whose accessibility decreased upon SOX9 degradation in a time-dependent manner (Figure 4D). We found that the loss of chromatin accessibility occurring at the G2/M border was progressively rescued in response to the restoration of SOX9 abundance upon PROTAC washout (Figure 4D, Supplementary Figure 3I, J). Other CREs showed a similar pattern of rapidly restoring the chromatin accessibility after an initial loss either in meta/anaphase (40 min.) or upon mitotic exit (120 min.) (Figure 4D). We also depicted some CREs whose chromatin accessibility decreased only in post-mitotic cells when SOX9 protein abundance was fully re-established, presumably caused by downstream effectors. We also identified CREs whose chromatin accessibility was increased upon SOX9 turnover (Supplementary Figure 4A). Nevertheless, the distal regions that resulted responsive to SOX9 perturbation were characterized by H3K4me1 and H3K27Ac histone marks and the binding of MED1, a subunit of the Mediator complex, thus representing putative enhancers (Figure 4E and Supplementary Figure 4B). However, the chromatin pattern of the SOX9-responsive CREs showed a lower H3K27Ac level with respect to the non-responsive putative enhancers, irrespective of the time point being analyzed (Figure 4E). Indeed, by grouping the SOX9-responsive CREs, we observed that they were characterized by reduced chromatin accessibility, a lower level of H3K27ac, and Mediator MED1 binding (Figure 4F, G). The reduced activation of the SOX9-responsive CREs was further confirmed by the lower expression level of the associated eRNAs depicted by nascent RNA-seq (Figure 4H). TF binding analyses showed enrichment for SOX motifs within the responsive CREs (Figure 4I, J, and Supplementary Figure 4C-E), which was not retrieved among CREs showing an increase of chromatin accessibility upon SOX9 depletion (Supplementary Figure 4F-G). This finding suggested a direct contribution of SOX9 to the maintenance of chromatin accessibility during mitosis. To verify this hypothesis, we investigated whether SOX9 chromatin occupancy correlated with the diminished chromatin accessibility occurring upon its degradation during mitosis. We found that SOX9 binds with relatively low affinity to SOX9-responsive CREs (Figure 4K and Supplementary Figure 4H), irrespective of the relative chromatin accessibility and nucleosome occupancy, or sequence biases (Supplementary Figure 5A-D). Indeed, nucleosome occupancy at the SOX9-responsive CREs was preserved throughout mitosis (Supplementary Figure 5E). However, we observed a time-dependent increase in nucleosome occupancy in response to SOX9 degradation, mirroring the transient reduction of chromatin accessibility (Supplementary Figure 5F). In sum, these results show that SOX9 contributes to maintaining chromatin accessibility at weak enhancers, throughout mitosis.

**Figure 4.**
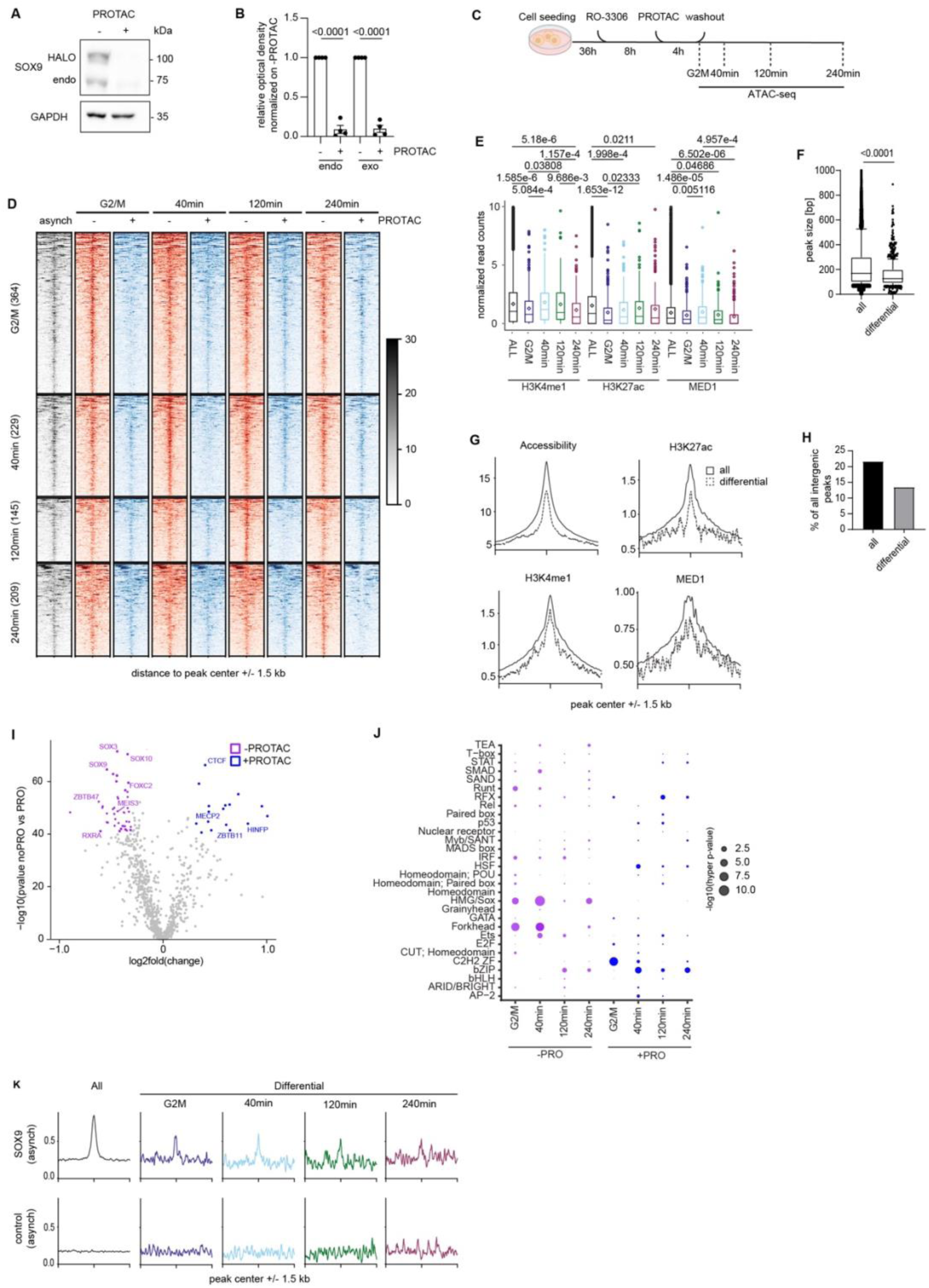
Acute depletion of SOX9 results in changes in chromatin accessibility dynamics and gene expression during mitosis. (A) Immunoblot of the endogenous SOX9 and exogenous SOX9-Halo without or after addition of 500nM Halo-PROTAC for 4 hours. Samples were analyzed at the asynchronous state. GAPDH was used as a loading control. (B) Quantification of SOX9 protein abundance retrieved by immunoblotting. Relative optical density was normalized on protein levels for endogenous SOX9 and exogenous SOX9-Halo without Halo-PROTAC3. P-values is determined using two-tailed t-test are reported in the Figure; n = 4 biological replicates. (C) Scheme showing the synchronization protocol to investigate the impact of SOX9 depletion on gene expression and chromatin accessibility during mitosis. Cells were synchronized 12 hours with the CDK1 inhibitor RO-3306 to arrest them at the G2/M border. 4 hours before the complete arrest, 500 nM Halo-PROTAC3 were added to deplete SOX9. Inhibitor and PROTAC were washed out and cells were released into mitosis; samples were taken at the indicated timepoints. (D) Heatmap of differential distal CREs showing a loss of chromatin accessibility upon SOX9 depletion at the timepoints G2/M border, 40 min, 120 min and 240 min post-release from G2/M arrest. (E) Boxplots showing the read counts for H3K4me1, H3K27ac and MED1 depicted by CUT&RUN on the CREs showing a loss of chromatin accessibility in asynchronous MD tumoroids upon SOX9 depletion, at the indicated timepoints. Significance was assessed by one-sided t-test. Significant (< 0.05) p-values are shown in the Figure. (F) Comparison of peak sizes of ATAC-seq retrieved at all the distal (ALL) and SOX9-responsive (differential) CREs. Shown are the mean, 10-90^th^ quartile as indicated by box borders. P-values defined by two-tailed unpaired t-test are indicated in the Figure. (G) Cumulative plots of peaks retrieved from all the distal (ALL) and SOX9-responsive (differential) CREs for chromatin accessibility, H3K4me1, H3K27ac and MED1 enrichment, defined by CUT&RUN in asynchronous cells. Differential peaks are shown as dashed lines, all peaks per cluster as continuous lines (H) Barplot showing the frequency of intergenic distal CRE with eRNA expression for either all peaks or peaks losing accessibility after SOX9 depletion. (I) TF Footprint analysis for different TF binding at distal *cis*-regulatory elements comparing the ATAC-seq signal at differential peaks in cells -/+ PROTAC at the G2/M border. Plotted are the log2fold change on the x-axis and the -log10(p-value) on the y-axis. TFs of interest are labeled. Full list of TFs is provided in Supplementary Table 5. (J) Analysis of relative TF family enrichment per timepoint comparing the -PROTAC and +PROTAC condition. Included were TF families with an overall number of members > 2. The -log10 of the hypergeometric t-test derived p-value is indicated by the dot size. (K) Cumulative plot of SOX9 chromatin occupancy depicted in asynchronous cells at distal CREs losing accessibility at the indicated timepoints. A control IgG is shown as control. Window shows peak center +/- 1.5 kb.

### MKM cooperates with SOX9 in mitotic bookmarking

Next, we thought of identifying the mechanism by which SOX9 maintains enhancer function during mitosis and which co-factors are the major players in helping to inherit memory to the daughter generation. To narrow down the possible interactors, we first checked for the interactors of SOX9, which may play a key role in supporting transcriptional memory. By querying the SOX9 interactors whose protein abundance was preserved during mitosis^49^, we identified a subset of chromatin factors that may cooperate with SOX9 in mitotic bookmarking (Figure 5A). As SOX9 has been shown to directly interact with members of the MKM such as MED12 and CDK8 (Supplementary Figure 6A)^50 51^, which specify the function of oncogenic enhancers in supporting transcriptional memory in metastatic cells^5^, we aimed at elucidating the role of MKM in mitotic bookmarking. To analyze the contribution of MED12, we analyzed its chromatin occupancy during mitosis by performing Cleavage Under Targets and Tagmentation (Cut&Tag) at the indicated time points (Figure 5B). By analyzing the MED12 genome distribution at distal CREs clustered based on the histone marks pattern (Supplementary Figure 6B, C), we found that MED12 was enriched at both active and poised putative enhancers (Supplementary Figure 6D). Of importance, its relative enrichment and genomic distribution resulted minimally perturbed upon entry into mitosis (Supplementary Figure 6D). To determine whether MED12 chromatin loading was depending at least in part on SOX9, we compared its genomic distribution and relative abundance on the SOX9-responsive CREs, in response to SOX9 degradation (Figure 5B, C and Supplementary Figure 6E). We found that SOX9-dependent CREs that are maintained accessible upon entry into mitosis and in metaphase were characterized by MED12 chromatin binding. Of importance, we observed that upon SOX9 degradation, MED12 association was strongly reduced during mitosis to then being fully rescued in the daughter cells when SOX9 protein level is fully re-established (Figure 5B, C, and Supplementary Figure 6E). This pattern was not depicted for CREs showing a SOX9 dependency during cytokinesis and post-mitotically, indicating a mitotic-specific interplay between SOX9 and MED12. To further decipher the contribution of SOX9-mediated loss of MED12 at distal sites, we separated the differential peaks obtained by ATAC-seq into peaks with or without a SOX TF motif (Figure 5D). Plotting MED12 distribution over these regions showed that MED12 is enriched at sites with a SOX motif, and its binding was reduced during mitosis in response to SOX9 degradation (Figure 5D). Of importance, this pattern was not retrieved when we considered distal CREs that showed an increment of accessibility in response to SOX9 perturbation (Figure 5C, d and Supplementary Figure 4A), further strengthening a direct contribution of MED12 in SOX9-mediated mitotic activity. These results suggest a cooperativity between SOX9 and MED12 in the mitotic bookmarking of metastatic cells.

**Figure 5.**
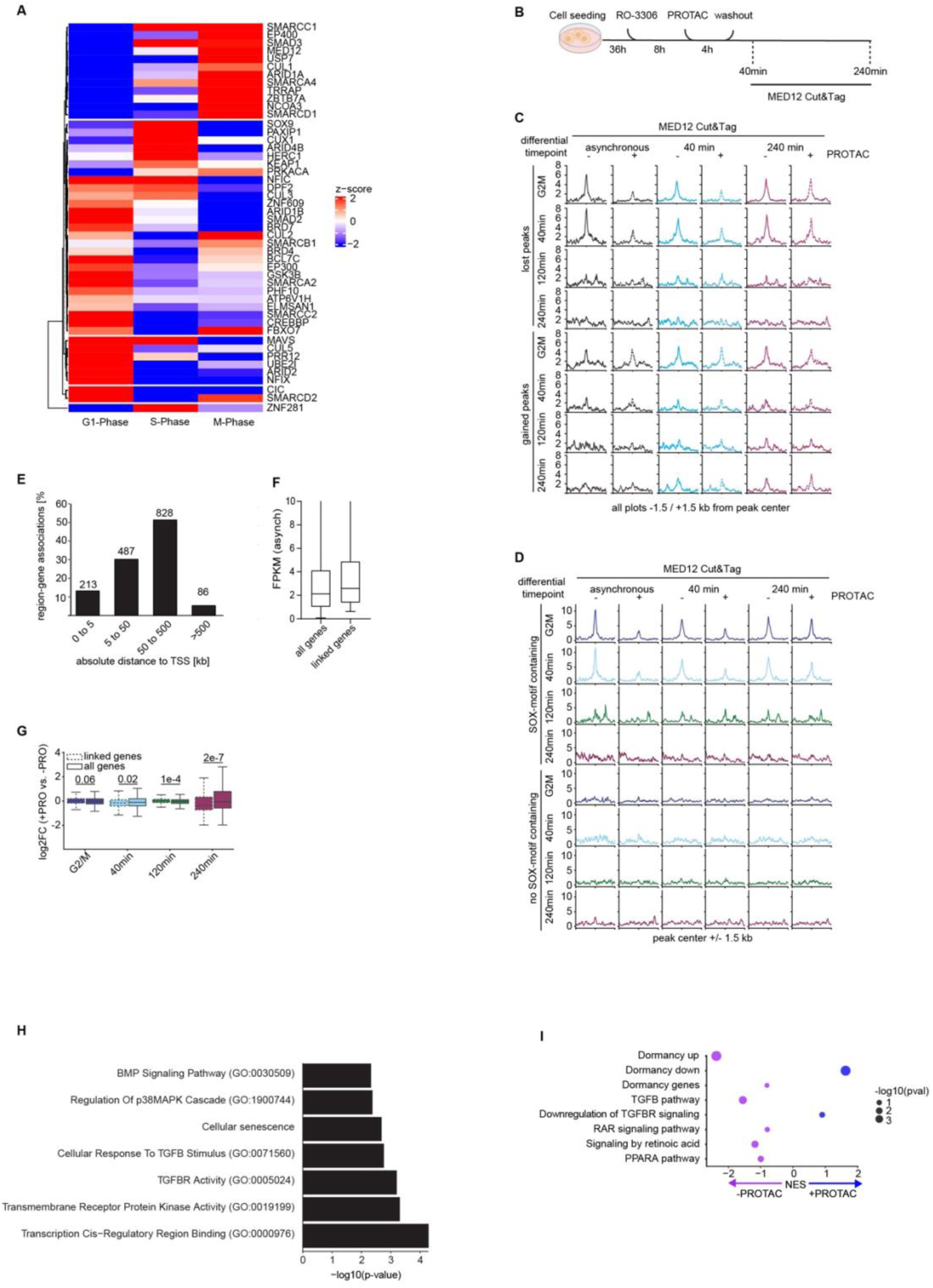
SOX9 depletion affects MED12 distribution during mitosis. (A) Heatmap depicting the cell cycle-dependent protein abundance of SOX9 interactors. Data are shown as z-score of the log2-normalized protein levels in G1-, S- and M-phase. (B) Scheme depicting the experimental setup for MED12 distribution by CUT&TAG after depletion of SOX9 prior to mitosis. (C) Cumulative plots of MED12 profiles at peaks either gaining or losing chromatin accessibility at the indicated timepoints, as determined by ATAC-seq. Shown is the MED12 signal centered on the peak center at the indicated timepoints. PROTAC samples are shown with dashed line, untreated samples with a continuous line. (D) Cumulative plots of MED12 profiles at distal CRE losing chromatin accessibility upon SOX9 depletion harboring a SOX9 TF DNA binding motif (top four rows) or not (bottom four rows). Shown is the MED12 signal centered on the peak center at the indicated timepoints. (E) Absolute distance in kilobases (kb) of peaks from the TSS of neighboring genes, as determined by GREAT. Percent of all peaks is indicated on the y-axis. (F) Comparison of gene expression levels (FPKM) of either all genes or genes being linked to differentially regulated distal CRE upon SOX9 depletion. (G) Comparison of log2fold changes of gene expression between -PROTAC and +PROTAC samples per timepoint for either all genes (full lines) or genes linked to differential timepoints (dashed lines). Shown are the mean, 10-90^th^ quartile as indicated by box borders and min/max values indicated by whiskers. P-values shown in the Figure were calculated using the Kolmogorov Smirnov test. (H) Pathway analysis of genes being linked to altered distal CRE upon SOX9 depletion. Shown is the -log10(adjusted p-value) of selected pathways. (I) Gene set enrichment analysis (GSEA) of genes expressed at 240 min after G2/M arrest washout upon SOX9 depletion (+PRO), or in the unperturbed condition (-PRO). The normalized enrichment score (NES) for selected gene sets is shown as values on the x-axis and -log10(p-value) is indicated by the dot size. GSEA was done by log2 ratio of classes with permutations of the gene set. For gene lists for the terms analyzed, see Supplementary Table 7.

### SOX9 primes the transcriptional reactivation of dormancy-specific genes

To rule out whether the SOX9-mediated mitotic bookmarking ensures the propagation of a metastatic-related transcriptional program to the daughter cells, we determined the changes in gene expression in response to SOX9 degradation in mitosis. We found that the perturbation of SOX9 abundance shortly before the G2/M border did not cause relevant changes in gene expression throughout mitosis, apart from post-mitotic genes showing a reduced transcript level with respect to the SOX9-unrelated genes, indicative of an improper gene expression reactivation (Figure 5E-G and Supplementary Figure 6F). To rule out the factors contributing to the modest responsiveness of genes linked to SOX9-responsive CREs during mitosis, we first examined their involvement in various biological functions. Functional pathway analyses showed enrichment for developmental and signal transduction pathways, including Wnt, BMP, TGFβ and p38 pathways, many of which are known regulators of cancer dormancy and reawakening^1,4,5,8–11^ (Figure 5H and Supplementary Figure 6G). To further verify whether SOX9 mitotic bookmarking contributed to the transcriptional reactivation of genes controlling cancer dormancy, we analyzed the changes in gene expression occurring in the daughter cells upon SOX9 degradation. GSEA analyses showed that dormancy-related signature and signaling pathways supporting quiescence were downregulated in response to SOX9 perturbation in mitosis (Figure 5I and Supplementary Figure 6H). Overall, these analyses showed that SOX9 mitotic bookmarking contributes to the transcriptional reactivation of dormancy-related genes in the daughter cells. Considering that signal-responsive TFs control the expression of the dormancy genes, it suggests that SOX9 primes the corresponding CREs during mitosis to ensure their transcriptional reactivation in response to the specifying signaling.

### SOX9 mitotic bookmarking favors entry into quiescence

Next, we verified whether SOX9 mitotic bookmarking primes for the transcriptional reactivation of a quiescent-related program, which entails daughter cells entering the G0 phase. To this end, we temporally arrested the *r*SOX9 metastatic tumoroids at the G2/M border, treated them with the HALO-PROTAC3 to degrade the HALO-SOX9 (Figure 4A), to then measure the frequency of quiescent, p27-positive (p27+) cells^5^ upon exiting from mitosis (Figure 6A). Time-course analyses showed that the frequency of p27-positive cells increased shortly after exiting mitosis to reach its highest level 8h after release from the G2/M border during the early G1 phase (Figure 6B, C). Of importance, SOX9 degradation before entry into prometaphase hampered the increment of p27-positive cells measured within the daughter cells (Figure 6B, C). Based on these findings, we asked whether the SOX9-dependent recruitment of the MKM module on the distal CREs during mitosis (Figure 5A-C) contributed to the priming of metastatic cells for entering quiescence after cell division. To address this point, we perturbed the kinase activity of the MKM upon entry into mitosis by transiently inhibiting CDK8 (Figure 6D). We found that the inhibition of the MKM kinase activity strongly reduced the frequency of p27-positive cells retrieved in the metastatic cells (Figure 6D-F), indicating a penetrant contribution to the reactivation of a quiescent state.

**Figure 6.**
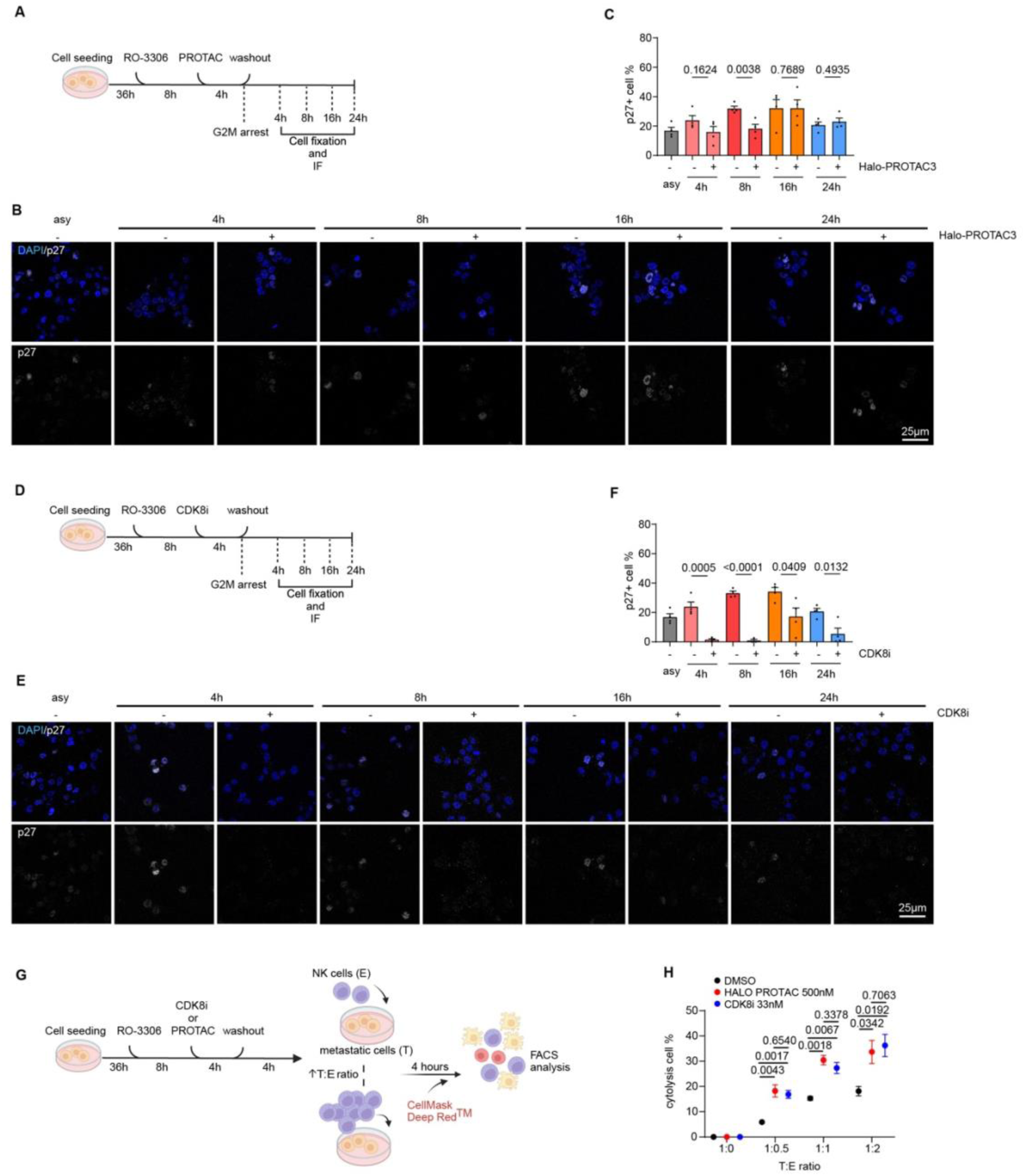
Loss of SOX9 prior to mitosis reduces quiescent population in the daughter generation and NK-cell mediated killing. (A) Scheme depicting the experimental setup for determining the effects of SOX9 depletion on the frequency of p27-positive cells after completion of mitosis. (B) Representative images of p27 immunostaining in cells treated with 500 nM Halo-PROTAC3 to deplete SOX9, as schematized in Figure 6A. DNA was labeled using DAPI. Scale bar: 25 µm. (C) Quantification of percentage of p27-positive cells after mitosis either with or without depletion of SOX9 before cell division. Data are means of three biological replicates +/- S.E.M. (n=3). Significance was determined by two-sided unpaired t-test. P-values are reported in the Figure. (D) Scheme depicting the experimental setup for determining the effects of CDK8 inhibition on the frequency of p27-positive cells after completion of mitosis. (E) Representative images of p27 immunostaining in cells treated with CDK8 inhibitor (CDK8i), as depicted in panel 6D. DNA was labeled using DAPI. Scale bar: 25 µm. (F) Quantification of percentage of p27-positive cells after mitosis either with or without inhibition of CDK8 before cell division. Data are means of three biological replicates +/- S.E.M. (n=3). Significance was determined by two-sided unpaired t-test. P-values are reported in the panel. (G) Schematic representation of the experimental setup for NK cytotoxicity assays on *r*SOX9 tumoroids after either depletion of SOX9 or inhibition of CDK8 before mitosis. (H) Average percentage of NK-mediated cytotoxicity of *r*SOX9 tumoroids after either depletion of SOX9 or inhibition of CDK8, respect to vehicle-only cells. Different ratios of target cells (T, cancer cells) or effector cells (E, NK cells) were used. Data are means of three biological replicates +/- S.E.M. (n=3). Significance was determined by two-sided unpaired t- test. P-values are reported in the panel.

Finally, we evaluated whether mitotic bookmarking increased the fitness of metastatic cells by preserving them from NK-mediated immunosurveillance^2–8^. Indeed, we have previously shown that entry into quiescence in response to pro-dormancy signals permits metastatic cells to escape from NK killing, thus fostering metastatic outbreaks^5^. To investigate whether SOX9/MKM priming of enhancers during mitosis increases the cellular fitness of metastatic tumoroids, we arrested the cells at the G2/M border and transiently depleted SOX9 or inhibited CDK8 (Figure 6G). Afterward, metastatic cells were co-cultured with different ratios of IL2-activated NK cells, and NK-mediated toxicity was determined. Upon depletion of SOX9 or inhibition of CDK8 we measured an increase in NK-mediated tumor cell killing compared to the untreated condition (Figure 6H). In sum, these results indicated that SOX9 mitotic bookmarking supported the reactivation of a dormancy-specific transcriptional program that contributed to entering quiescence, thus facilitating the escape from NK surveillance.

## DISCUSSION

To correctly divide and transmit their identity to the progeny, cells must preserve the chromatin state during mitosis and reactivate the transcriptional program upon cell division. We aimed to identify the pro-metastatic TFs important for establishing this permissive state in metastatic cells. We specifically addressed whether the priming of enhancers during mitosis contributed to metastatic cell plasticity exemplified by their capability to either progress in the cell cycle or enter quiescence shortly after cell division. By analyzing the changes in chromatin accessibility occurring along mitosis, we found that besides promoters, some of the distal CREs also preserved an accessible chromatin state. When analyzing TF footprint changes during mitosis, we found that subsets of TF motifs were specifically enriched at putative enhancers that were maintained accessible during prometaphase. This finding differs from previously published results, which showed that mainly the TSS remain in an open and bookmarked state, whereas we also detected a similar chromatin pattern at distal CREs^19^. These apparent differences may depend on the chosen time window, as we analyzed for the first-time chromatin accessibility changes occurring early in prometaphase, which showed a trajectory of closing and opening, thus not representing a canonical “bookmarking” event. Among the identified TFs bookmarking the enhancers during the early stages of mitosis, we identified SOX9, a pro-metastatic TF, which is often deregulated in breast cancer^5,11, 52^. To better understand of the potential mechanisms by which TFs could contribute to priming enhancers for their reactivation upon cell division, we first analyzed their gene expression pattern during mitosis. We found that the expression of most TFs was markedly reduced upon entry into mitosis, followed by a wave of reactivation at different time points. However, we identified a small cluster of TFs, among which SOX9 and other cancer-associated TFs, whose expression was marginally reduced upon entry into mitosis and was further induced during ana/telophase. This peculiar gene expression pattern suggests that these TFs may play a key role in the mitotic bookmarking of metastatic cells. Indeed, to maintain cell identity while preserving cell plasticity, post-mitotic gene expression reactivation and enhancer priming must be tightly coordinated to successfully propagate the transcriptional program and the responsiveness to specific signals^5,23^. Of importance, perturbation of TF function during mitosis can alter the cellular outcome^24^, linking single TF activity to the maintenance of cellular phenotypes. SOX9 has been described as a master regulator of cell fate in different tissues, making it a likely candidate for driving memory propagation through mitosis. To better understand the function of SOX9 in this context, we first analyzed SOX9 distribution during mitosis, which revealed a partial retainment of SOX9 on mitotic chromatin, comparable to the kinetics of KLF4 and FOXO1^28^. This partial persistence of SOX9 on mitotic chromatin might implicate two different aspects: a) SOX9 is important for bookmarking and, b) its activity in prometaphase is relevant to establish a chromatin state for transferring information to the progeny. To tackle this, we employed targeted degradation of SOX9 in a timely manner using a PROTAC-mediated approach. Interestingly, compared to other targeted TFs^53^, SOX9 expression was rapidly restored after the washout of the PROTAC, thus permitting to specifically determine its contribution during prometaphase. Our data show that the loss of SOX9 prior to mitosis reduces chromatin accessibility at certain distal CREs, whose accessible chromatin state was regained at later stages of mitosis once SOX9 protein level was re-established. These findings align with SOX9 being described as a potential pioneering factor, capable of reprogramming the epigenetic states of healthy and tumorigenic cells^5,54,55^. Interestingly, the SOX9-responsive CREs resulted in primed enhancers at which SOX9 binds with relatively low affinity. This chromatin pattern may explain their responsiveness to the time-restricted perturbation of SOX9 in prometaphase, and the resulting modest changes in the gene expression pattern of the linked genes upon mitotic exit. Indeed, we found that the putative-regulated genes are enriched for signal transduction pathways involved in supporting cancer dormancy, including p38 MAPK cascade, RAR, TGFβ, and BMP signaling pathways^1,4,5,8^. To gain biological insights into the possible mechanism by which SOX9 primed these signal-responsive enhancers during mitosis, we queried its interactome and found that its binding is functional for the recruitment of the MKM. Besides modulating SOX9 activity, recent findings highlight the contribution of the MKM in tuning transcriptional responses to signaling cascades in different contexts by modulating enhancers function^36,41–44^. Albeit these evidence, we reported for the first time the contribution of MKM to mitotic bookmarking to preserve the responsiveness of metastatic cells to pro-dormancy signals. Indeed, we showed that either the mitotic perturbation of SOX9 or MKM activity impinged the capability of metastatic cells to enter quiescence upon cell division. Furthermore, as this state protects from NK-mediated immune surveillance^5,7^, we tested the contribution of SOX9-mediated enhancer priming during early mitosis to escape NK killing. We found that while SOX9 mitotic bookmarking was necessary to preserve the capability of metastatic cells to evade NK surveillance, MKM inhibition showed a more remarkable effect, suggesting that it can cooperate with other pro-dormancy TFs. These findings are highly relevant, considering that genes coding for MKM components are frequently mutated in cancer, yet their contribution to immune evasion has not been tested so far. Besides MKM module, we also identify other SOX9 cofactors that could potentially contribute to is bookmarking activities. In particular, we retrieved many chromatin factors belonging to the SWI/SNF remodeling complex, which have been recently shown to contribute itself to mitotic bookmarking^33^. Whether the interplay between SOX9, MKM and SWI/SNF complex is necessary to execute its mitotic bookmarking in the context of metastatic cells remains an open question that requires further studies. With this study, we showed that the interference with the activity of a pro-metastatic TF prior to and during early mitosis is sufficient to alter the cell choice between proliferation and quiescence. This finding greatly impacts for the development of novel therapeutic approaches targeting the molecular mechanisms controlling cancer dormancy.

## DATA AVAIABILITY

The next generation sequencing datasets (ATAC-seq, nascentRNA-seq and Cut&Tag) reported in this paper will be uploaded to GEO. Data will be made publicly available once the manuscript is accepted.

## AUTHOR CONTRIBUTION

A.Z. and S.B. conceived the study, designed the experiments, and interpreted the data. S.B., D.M., L. F., M.L.N., and E.F. performed the cellular and molecular biology studies, and participated in data analyses.; NGS data analyses were performed by S.B. and L.F. A.Z. and S.B. wrote the manuscript.

## ACKNOLEDGMENT

We are grateful for the help of the team of the CIBIO NGS facility, Dr. Veronica De Sanctis, Dr. Roberto Bertorelli and Dr. Paolo Cavallerio, for their helpful contribution and sample sequencing, the CIBIO imaging facility for help with microscopy. We thank Prof. Luca Fava for the scientific discussion technical suggestions related to mitotic cell synchronization, and for sharing reagents. We also thank all the current and past and present members of the Zippo Lab for helpful advice, assistance, and carefully reading the manuscript.

