## Supplementary Data for "Metastatic cell plasticity is maintained by SOX9 function during mitosis in breast cancer"

### Supplementary Figures

#### Supplementary Figure 1

**(A)** FACS plot showing mitotic arrest efficiency using RO-3306 and subsequent Nocodazole incubation in metastasis-derived cells (MD). Cells were stained using Propidium iodide (x-axis) to determine DNA content and pH3-Ser10 (y-axis) as marker of prometaphase. **(B)** Barplots showing percentage of genome-wide distribution of chromatin accessible regions in XD and MD tumoroids. Categories shown are 3' untranslated region (UTR), 5'UTR, Exon, Intergenic, Intron, mRNA Promoter/TSS (-1000 bp/+100 bp from TSS) and transcription termination site (TTS). The log2-transformed enrichment score for observed/expected per category is shown on the right. **(C)** Heatmaps (left) and profiles (right) showing chromatin accessibility determined with ATAC-seq at distal *cis*-regulatory elements (CREs) (top) and TSS peaks (bottom) peaks in XD cells in the asynchronous condition, during early and late mitosis, as indicated in the scheme. Signal is centered on the peak center with a window of -/- 2.5 kb. Shown is the intensity of two merged replicates. **(D)** Heatmaps and profiles showing changes in chromatin accessibility at the timepoints asynchronous, early and late prometaphase, at distal CREs showing motifs for the indicated TFs, SOX9 (2085 peaks), JUNB (6965 peaks), and CTCF (6086 peaks), as determined by TFs footprints. **(E)** TF footprints analysis of distal CRE peaks comparing asynchronous vs. early prometaphase (top) and early vs. late prometaphase (bottom) in XD cells. Fold change is shown on x-axis, -log10(p-value) is shown on y-axis. Significant TF motifs for all three timepoints are colored and selected TFs are labeled. For full list of TFs, see Supplementary Table 2. **(F)** TF footprints analysis of TSS peaks comparing asynchronous vs. early prometaphase (top) and early vs. late prometaphase (bottom) in XD tumoroids. Fold change is shown on x-axis, -log10(p-value) is shown on y-axis. Significant TF motifs for all three timepoints are colored and selected TFs are labeled. For full list of TFs, see Supplementary Table 2. **(G)** TF footprints analysis of TSS peaks comparing asynchronous vs. early prometaphase (top) and early vs. late prometaphase (bottom) in MD tumoroids. Fold change is shown on x-axis, -log10(p-value) is shown on y-axis. Significant TF motifs for all three timepoints are colored and selected TF are labeled. For full list of TFs, see Supplementary Table 2.

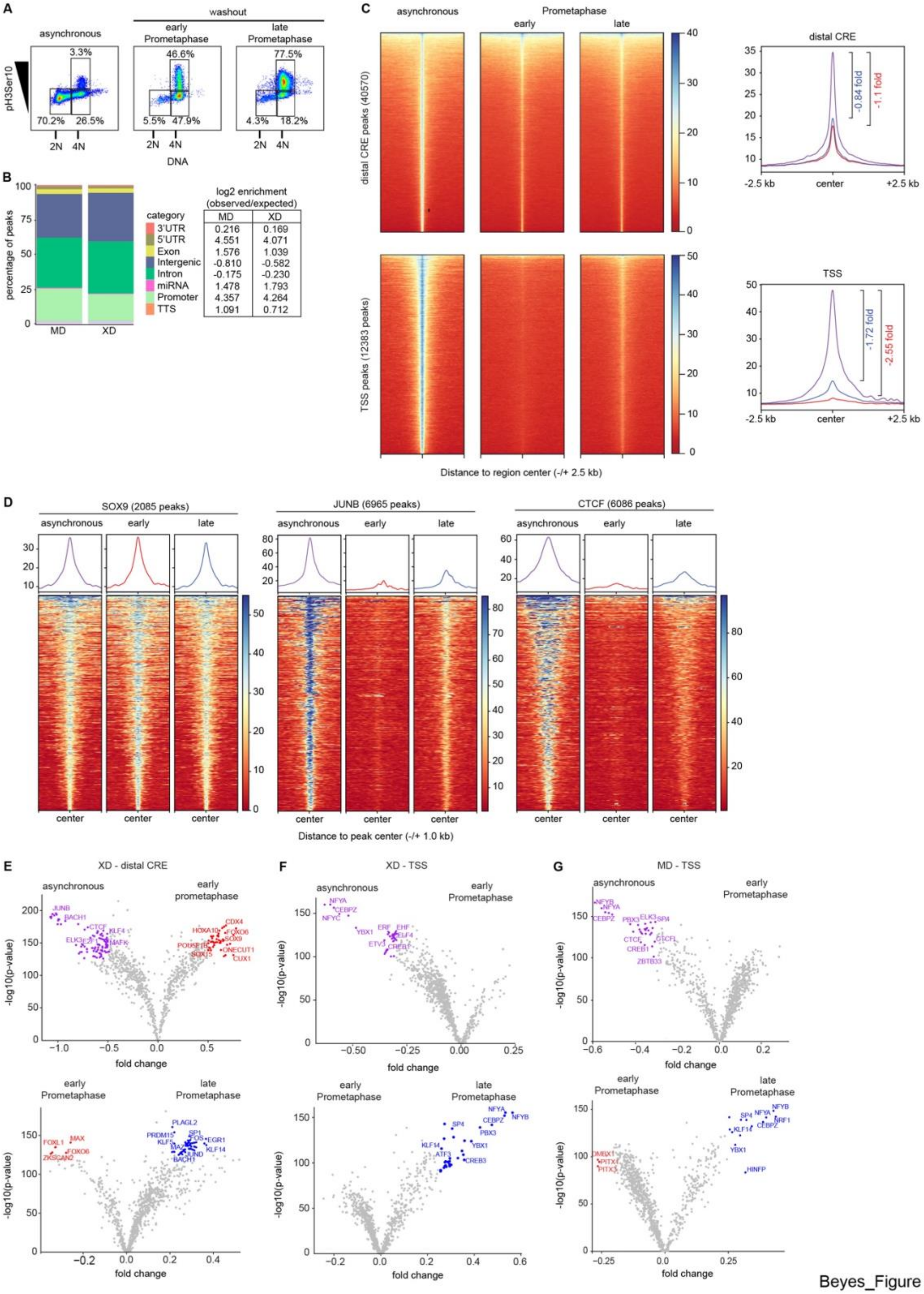


Beyes_Figure S1

**Supplementary Figure 2**

**(A)** Heatmap showing k-means clustering in 10 clusters of gene expression changes during mitosis after a G2/M arrest using RO-3306, as indicated in Figure 2a. Shown is the z-score of the FPKM values over the indicated timepoints per gene. For genes per cluster, see Supplementary Table 3. **(B)** Boxplot showing the z-score of the FPKM for each timepoint of reactivation for all timepoints analyzed. Black = asynchronous, dark blue = G2/M, light blue = 40min, green = 120min and violet = 240min. The number of genes per timepoint are indicated below in brackets. Timepoint of reactivation is based on the heatmap in panel S2A. **(C)** Pathways enriched for genes showing the first spike in expression at the G2/M arrest, selected terms from the top hits are shown. Y-axis shows -log10(adjusted.p-value). For full list of pathways, see Supplementary Table 3. **(D)** Pathways enriched for genes showing the first spike in expression after 40 minutes of release from G2/M arrest, selected terms from the top hits are shown. Y-axis shows -log10(adjusted.p-value). For full list of pathways, see Supplementary Table 3. **(E)** Pathways enriched for genes showing the first spike in expression after 120 minutes of release from G2/M arrest, selected terms from the top hits are shown. Y-axis shows -log10(adjusted.p-value). For full list of pathways, see Supplementary Table 3. **(F)** Pathways enriched for genes showing the first spike in expression after 240 minutes of release from G2/M arrest, selected terms from the top hits are shown. Y-axis shows -log10(adjusted.p-value). For full list of pathways, see Supplementary Table 3. **(G)** Browser view of LMNA, example gene for reactivation of expression after 40 min of release from G2/M arrest. Shown are the forward and reverse strand per timepoint asynchronous, G2/M border, 40min, 120min and 240min release. Coordinates shown: LMNA chr1:156,082,465-156,112,911. All coordinates are hg19. **(H)**

Immunoblot showing SOX9, CTCF and MYC protein levels in asynchronous cells, G2/M arrested cells by using RO-3306 and cells released from RO-3306 arrest at 40 / 80 / 120 / 160 / 200 /240 minutes. GAPDH was used as a loading control. **(I)** Quantification of protein levels during washout from RO-3306 mediated G2/M arrest. Intensity was normalized on the loading control (GAPDH) in three independent biological replicates. P-values determined by two-tailed unpaired t-test. p-values are reported in the figure.

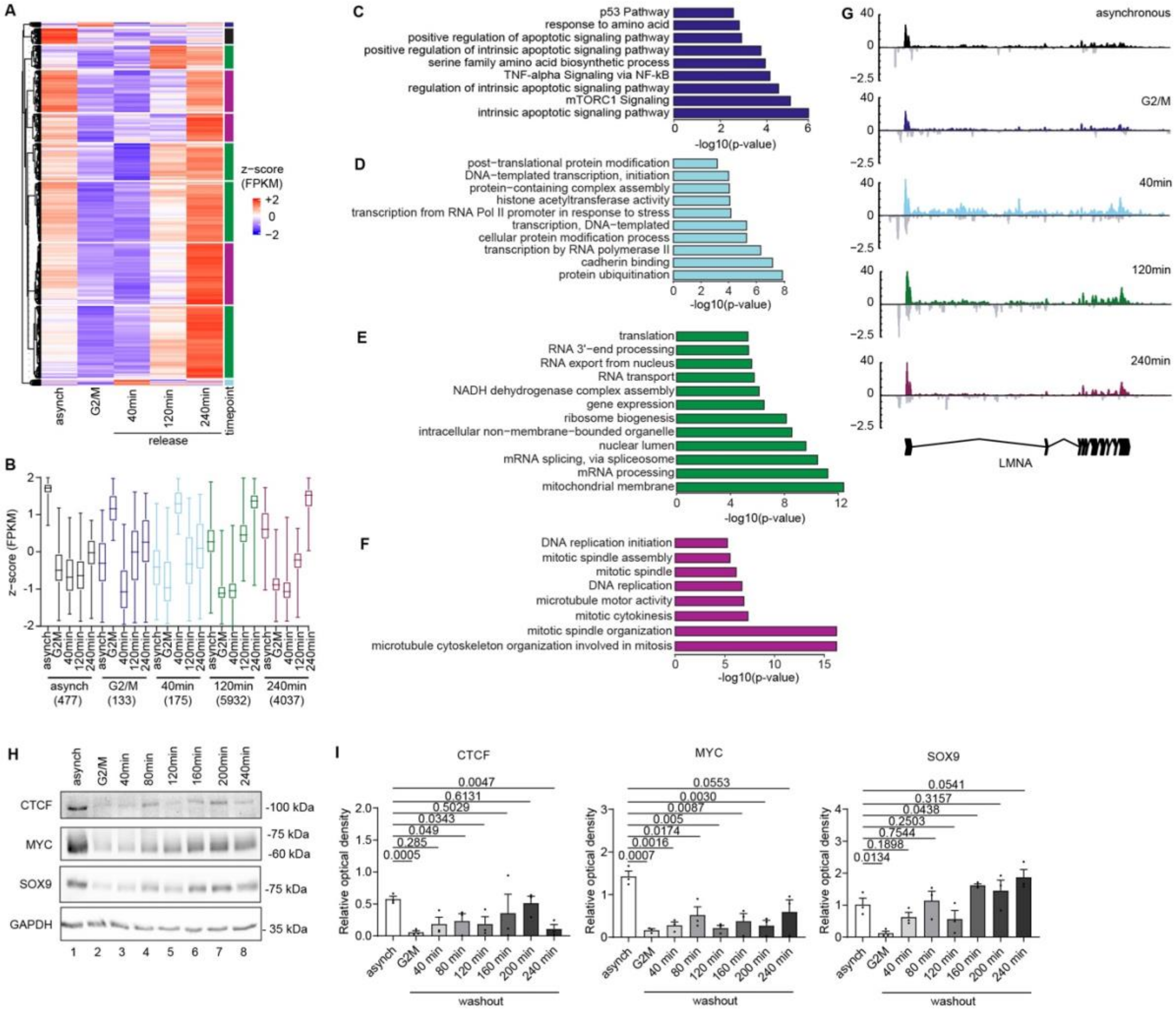
A
z−score
(FPKM)
+2
0
−2
asynch
G2/M
40min
120min
240min
timepoint
release
B
z−score (FPKM)
asynch (477)
G2/M (133)
40min (175)
120min (5932)
240min (4037)
C
p53 Pathway
response to amino acid
positive regulation of apoptotic signaling pathway
positive regulation of intrinsic apoptotic signaling pathway
serine family amino acid biosynthetic process
TNF-alpha Signaling via NF-kB
regulation of intrinsic apoptotic signaling pathway
mTORC1 Signaling
intrinsic apoptotic signaling pathway
-log10(p-value)
D
post−translational protein modification
DNA−templated transcription, initiation
protein−containing complex assembly
histone acetyltransferase activity
transcription from RNA Pol II promoter in response to stress
transcription, DNA−templated
cellular protein modification process
transcription by RNA polymerase II
cadherin binding
protein ubiquitination
-log10(p-value)
E
translation
RNA 3'−end processing
RNA export from nucleus
RNA transport
NADH dehydrogenase complex assembly
gene expression
ribosome biogenesis
intracellular non−membrane−bounded organelle
nuclear lumen
mRNA splicing, via spliceosome
mRNA processing
mitochondrial membrane
-log10(p-value)
F
DNA replication initiation
mitotic spindle assembly
mitotic spindle
DNA replication
microtubule motor activity
mitotic cytokinesis
mitotic spindle organization
microtubule cytoskeleton organization involved in mitosis
-log10(p-value)
G
asynchronous
G2/M
40min
120min
240min
LMNA
H
asynch
G2/M
40min
80min
120min
160min
200min
240min
CTCF
-100 kDa
MYC
-75 kDa
-60 kDa
SOX9
-75 kDa
GAPDH
- 35 kDa
1 2 3 4 5 6 7 8
I
CTCF
0.0047
0.6131
0.5029
0.0343
0.049
0.285
0.0005
MYC
0.0553
0.0030
0.0087
0.005
0.0174
0.0016
0.0007
SOX9
0.0541
0.3157
0.0438
0.2503
0.7544
0.1898
0.0134
Relative optical density
asynch
G2M
40 min
80 min
120 min
160 min
200 min
240 min
washout

**Supplementary Figure 3**

**(A)** Barplot indicating percentage of cells being positive of SOX9 in parental MD and *r*SOX9 tumoroids. Data were retrieved from three independent biological samples and P-value was determined by two-tailed unpaired t-test and is reported in the figure. **(B)** Violin plot showing nuclear mean intensity for SOX9 in parental and *r*SOX9 tumoroids. Data were retrieved from three independent biological samples and P-value was determined by two-tailed unpaired t-test and is reported in the figure. **(C)** Barplot indicating percentage of p27-positive cells in parental and *r*SOX9 tumoroids. Data were retrieved from four independent biological samples and P-value was determined by two-tailed unpaired t-test and are reported in the figure. **(D)** Violin plot showing nuclear mean intensity for p27 in parental and *r*SOX9 tumoroids. Data were retrieved from three independent biological samples and P-value was determined by two-tailed unpaired t-test and are reported in the figure. **(E)** Barplot indicating percentage of cells being positive of Ki67 in parental and *r*SOX9 tumoroids. Data were retrieved from three independent biological samples and P-value was determined by two-tailed unpaired t-test and is reported in the figure. **(F)** Violin plot showing nuclear mean intensity for Ki67 in parental and *r*SOX9 tumoroids. P-value was determined by two-tailed unpaired t-test and is reported in the figure. **(G)** Dotplot showing the correlation of the FPKM values between the expressed genes of MD and *r*SOX9 tumoroids by nascent RNA-seq in the asynchronous state. Correlation coefficient is reported in the figure. **(H)** Quantification of SOX9 HALO signal intensity at different cell cycle stages as shown in Figure 3D. P-values determined by two-tailed unpaired t-test. Exact p-values are reported in the figure. **(I)** Immunoblot of SOX9 protein levels retrieved in *r*SOX9 tumoroids upon release from G2/M arrest by RO-3306, in presence or absence of PROTAC3 treatment.

Protein levels for endogenous SOX9 levels and SOX9-Halo levels are shown. GAPDH was used as loading control. **(J)** Quantification of endogenous, exogenous and total SOX9 protein levels. Intensity was normalized on the loading control (GAPDH) and additionally calculated as fraction of the asynchronous -PROTAC condition. Shown are the mean +/- s.e.m of three biological replicates. Significance was calculated using an unpaired two-tailed t-test.

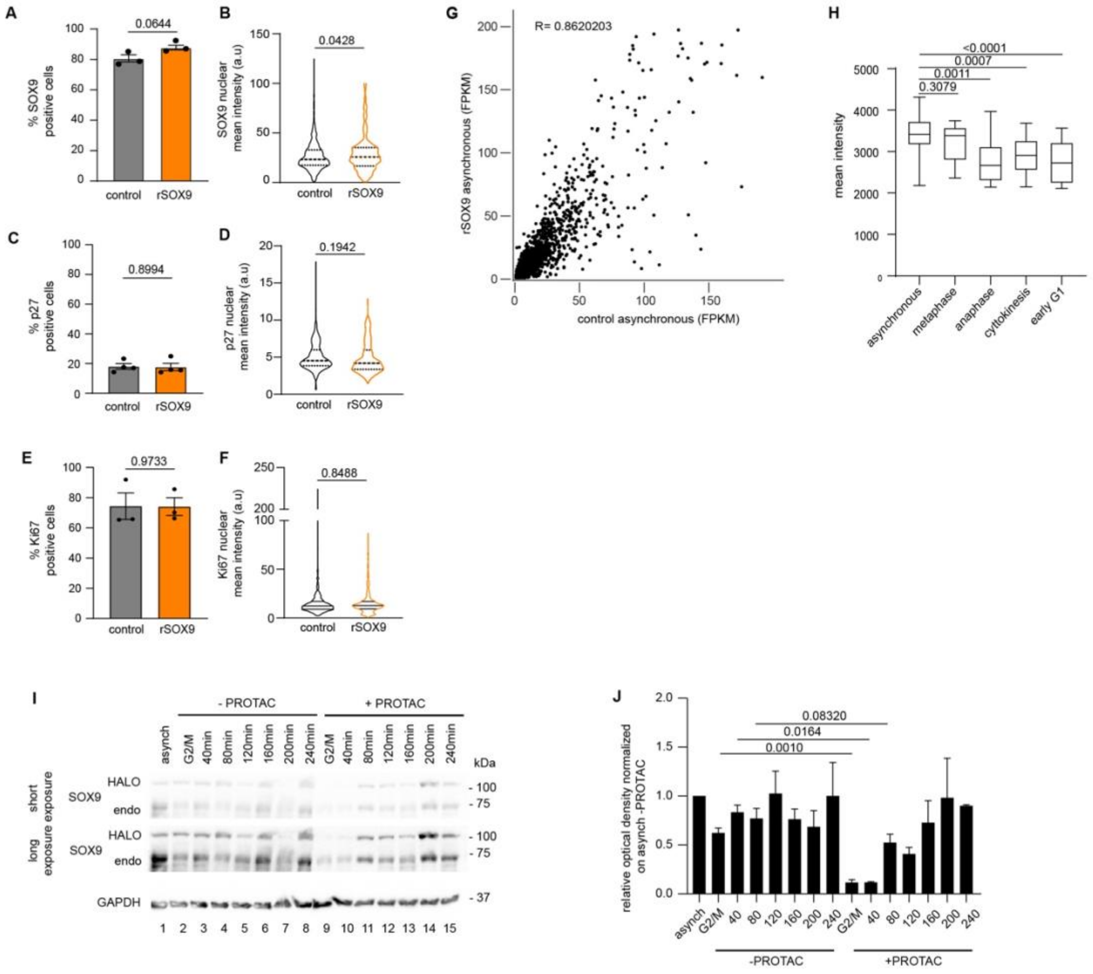


Beyes_Figure S3

**Supplementary Figure 4**

**(A)** Heatmaps of differential distal CREs showing an increase of chromatin accessibility upon SOX9 depletion at the timepoints G2/M border, 40 min, 120 min and 240 min post-release from G2/M arrest as indicated in Figure 4c). Shown is the signal +/- 1.5 kb from the peak center. **(B)** Boxplots showing the read counts for H3K4me1, H3K27ac and MED1 depicted by CUT&RUN on the CREs losing chromatin accessibility upon SOX9 depletion in asynchronous MD tumoroids. Timepoint of chromatin accessibility changes upon SOX9 depletion is color coded. **(C)** Volcano plot showing the TF footprint analysis for differential TF binding at CREs losing chromatin accessibility upon acute depletion of SOX9. Shown are the TF motifs enriched at 40min -/+ Halo-PROTAC3 post washout. Plotted are the log2fold change on the x-axis and the -log10(p-value) on the y-axis. Selected TFs are indicated. For full list of TFs, see Supplementary Table 5. **(D)** Volcano plot showing the TF footprint analysis for differential TFs binding at distal CRE*s* losing chromatin accessibility upon acute depletion of SOX9. Shown are the TF motifs enriched at 120min -/+ Halo-PROTAC3 post washout. Plotted are the log2fold change on the x-axis and the -log10(p-value) on the y-axis. Selected TFs are indicated. For full list of TFs, see Supplementary Table 5. **(E)** Volcano plot showing the TF footprint analysis for differential TF binding at distal CREs losing chromatin accessibility upon acute depletion of SOX9. Shown are the TF motifs enriched at 240min -/+ Halo-PROTAC3 post washout. Plotted are the log2fold change on the x-axis and the -log10(p-value) on the y-axis. Selected TFs are indicated. For full list of TFs, see Supplementary Table 5. **(F)** Volcano plot showing the TF footprint analysis for differential TF binding distal CREs gaining chromatin accessibility upon acute depletion of SOX9. Shown are the TF motifs enriched at G2/M (top left), 40min (top right), 120min (bottom left) and 240min (bottom right) after release from G2/M arrest and -/+ Halo-PROTAC3. Plotted are the log2fold change on the x-axis and the -log10(p-value) on the y-axis. Selected TFs are indicated. For full list of TFs, see Supplementary Table 6. **(G)** Analysis of relative TF family enrichment per timepoint comparing the -PROTAC and +PROTAC condition at peaks gaining chromatin accessibility upon SOX9 depletion. Included were TF families with an overall number of members > 2. The -log10 of the hypergeometric t-test derived p-value is indicated by the dot size. **(H)** Table showing number of peaks losing chromatin accessibility upon SOX9 depletion divided per quartile and timepoint of loss.

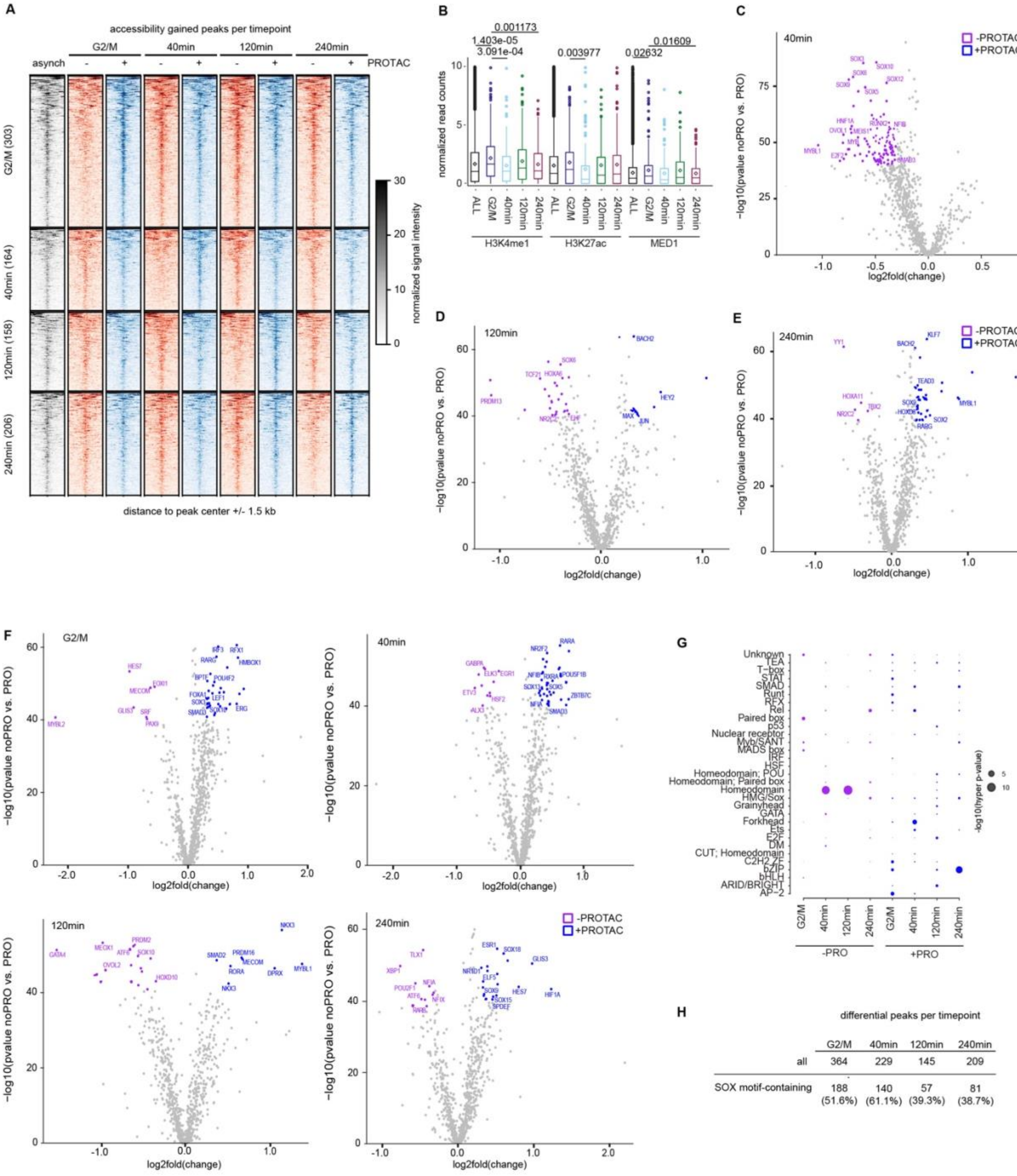


differential peaks per timepoint

| | G2/M | 40min | 120min | 240min |
|---|---|---|---|---|
| all | 364 | 229 | 145 | 209 |
| SOX motif-containing | 188 (51.6%) | 140 (61.1%) | 57 (39.3%) | 81 (38.7%) |

Beyes_Figure S4

**Supplementary Figure 5**

**(A)** Cumulative plot and heatmap of SOX9 chromatin binding on distal CREs containing SOX TF motifs, depicted in asynchronous MD tumoroids. Data were divided and represented in quartiles. **(B)** Cumulative plot and heatmap of ATAC-seq signal in asynchronous MD tumoroids on same regions as in panel A. **(C)** Nucleosome positioning analysis at either all peaks (continuous line) or SOX9-responsive CREs -differential peaks- (dashed line), identified by ATAC-seq per cluster of SOX9 enrichment, as determined in panel A. Shown is the signal centered on the peak center -/+ 0.5 kb. **(D)** Analysis of enrichment of repetitive elements in SOX9-responsive CREs. Shown are the observed distribution (red line) versus the distribution of 1000x permutated sequences (blue columns) of the same peak size. **(E)** Nucleosome positioning analysis of either all differential peaks, or differential peaks divided by timepoint in asynchronous cells. **(F)** Nucleosome positioning at peaks losing chromatin accessibility centered on SOX TF motifs at different timepoints analyzed either with or without depletion of SOX9 as indicated in Figure 4A.

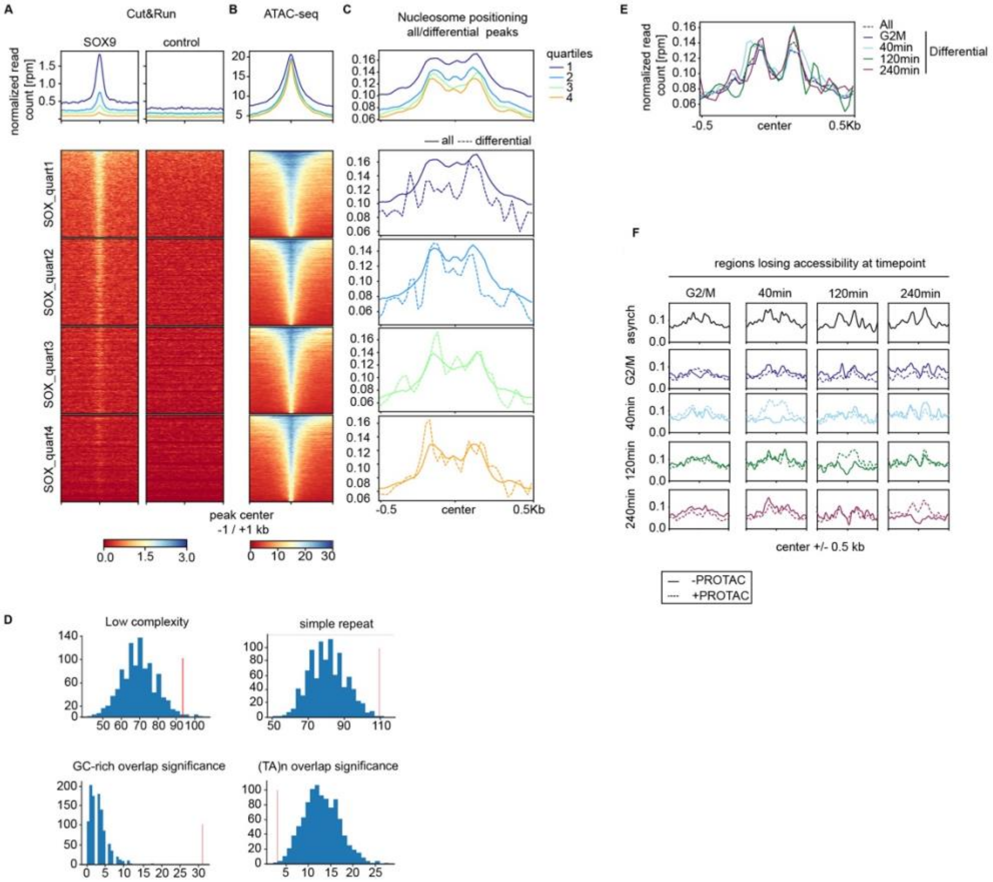

A
Cut&Run
SOX9
control
normalized read count [rpm]
B
ATAC-seq
C
Nucleosome positioning all/differential peaks
quartiles
1
2
3
4
— all ---- differential
SOX_quart1
SOX_quart2
SOX_quart3
SOX_quart4
peak center -1 / +1 kb
-0.5
center
0.5Kb
E
All
G2M
40min
120min
240min
Differential
F
regions losing accessibility at timepoint
G2/M
40min
120min
240min
asynch
center +/- 0.5 kb
-PROTAC
+PROTAC
D
Low complexity
simple repeat
GC-rich overlap significance
(TA)n overlap significance

**Supplementary Figure 6**

**(A)** Heatmap depicting the cell cycle-dependent protein expression levels of subunits of the Mediator complex. Shown is the z-score of the log2-normalized protein levels in G1-, S- and M-phase. Category indicates Mediator core unit or Mediator Kinase Module (MKM). **(B)** Heatmap of the ATAC-seq, H3K4me1, H3K27ac and MED1 signals retrieved in the asynchronous state. Overlapping peaks were divided into highly active regions (Accessibility + H3K4me1 + H3K27a), poised (Accessibility + H3K4me1) and regions showing only accessibility, but no overlap with active enhancer marks. Distance to peak center is +/- 1.5 kb. **(C)** Chromatin accessibility profiles determined by ATAC-seq, H3K4me1, H3K27ac and MED1 for the three clusters, defined in panel A. Profile plots are centered on the center of the overlap between the samples and shown is the profile +/- 1.5 kb. **(D)** Profiles (top) and heatmaps (bottom) of MED12 enrichment determined by CUT&TAG at all distal CRE peaks of the three different clusters determined by chromatin accessibility (ATAC-seq), H3K4me1 and H3K27ac peak distribution. As control, the IgG in the asynchronous setting is shown. **(E)** Principal Component Analysis (PCA) of MED12 signal determined by CUT&TAG for asynchronous, 40min, and 240min samples with or without depletion of SOX9 prior to mitosis (as depicted in Figure 5A) at SOX9-responsive CREs. **(F)** Boxplot showing the log2fold of gene expression for either all genes expressed at the indicated timepoint (left) or for genes with a log2fold > 0.5. Numbers are indicated in the table (top right). **(G)** Pathway analysis of genes being linked to SOX9-responsive CREs. Shown is the -log10(p-value) of selected pathways. **(H)** GSEA plots for dormancy signatures for genes being expressed at 240min, with or without SOX9 depletion. Graphs correspond to Figure 5I.

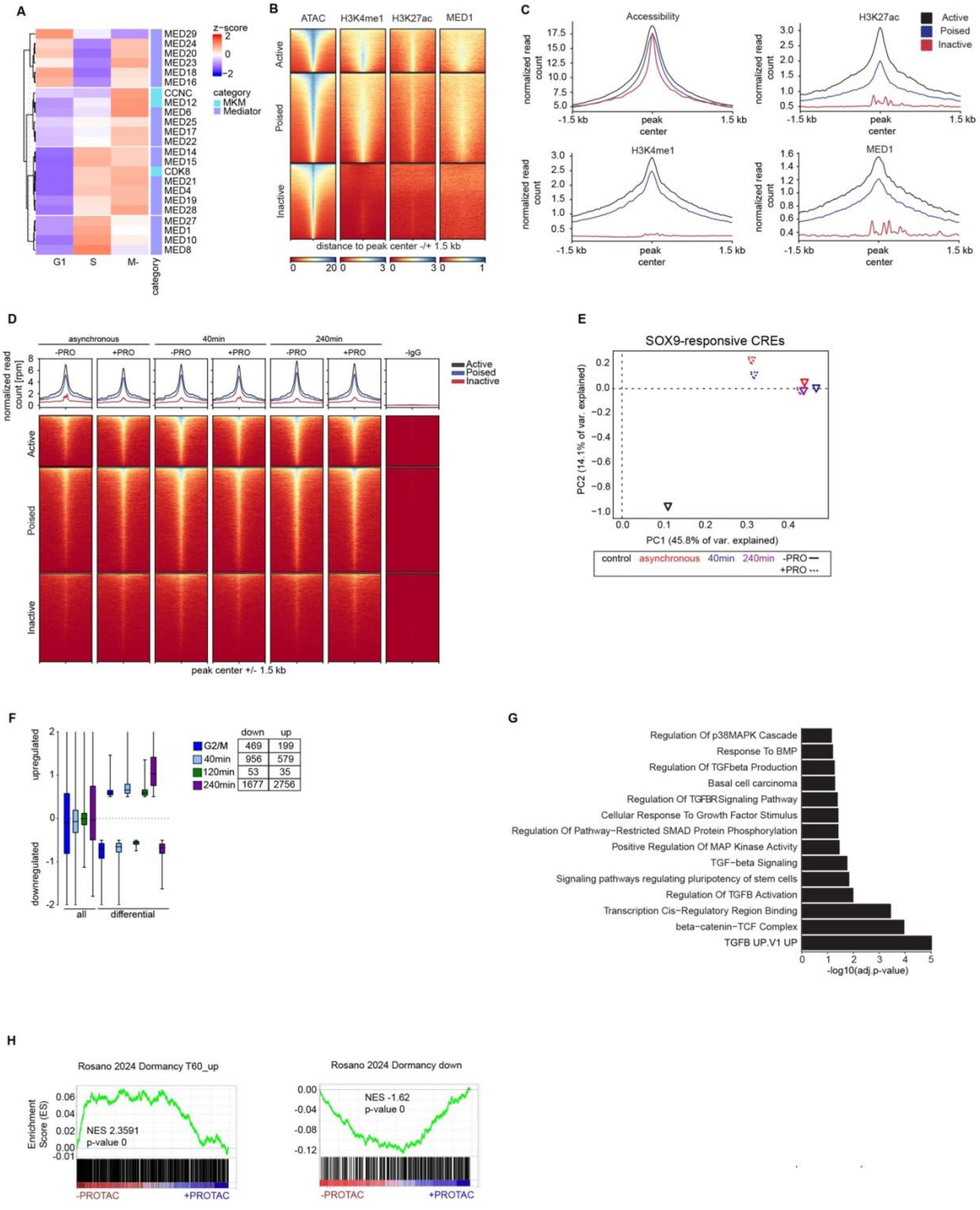


Beyes_Figure S6

#### Supplementary Tables

**Supplementary Table 1. TF footprints analysis at distal CREs in MD tumoroids at the timepoints asynchronous, early and late prometaphase; related to Figure 1.**

The table reports the results from the TOBIAS analysis in asynchronous, early and late prometaphase MD tumoroids.

**Supplementary Table 2. TF footprints analysis at distal CREs in XD cells, at TSS in XD and MD tumoroids at the timepoints asynchronous, early, and late prometaphase; related to Supplementary Figure 1.**

The table is separated into three sheets: XD_distal reports the results from the TOBIAS analysis at distal CREs in asynchronous, early and late prometaphase XD tumoroids. MD_TSS the results from the TOBIAS analysis at TSS in asynchronous, early, and late prometaphase MD tumoroids. XD_TSS the results from the TOBIAS analysis at TSS in asynchronous, early, and late prometaphase XD tumoroids.

**Supplementary Table 3. Nascent RNA-seq for gene expression reactivation after release from G2/M arrest; related to Supplementary Figure 2**

The table is divided into five sheets: Z-score reports the z-score for expressed genes in asynchronous, G2/M arrested and 40min, 120min and 240min released MD cells. Gene_cluster reports the k-means clustering of genes for their reactivation kinetics. GO_G2/M reports the functional pathways linked to the genes being reactivated at the G2/M border. GO_40min reports the functional pathways linked to the genes being reactivated after 40min of release. GO_120min reports the functional pathways linked to the genes being reactivated after 120min of release. GO_240min reports the functional pathways linked to the genes being reactivated after 240min.

**Supplementary Table 4. Clusters of TF gene expression reactivation after release from G2/M arrest; related to Figure 2**

The table reports the k-means clustering of TF gene expression after release from G2/M arrest.

**Supplementary Table 5. TF footprints analysis of distal CREs losing chromatin accessibility upon acute SOX9 depletion; related to Figure 4**

The table reports the results from TOBIAS analysis at distal CREs losing chromatin accessibility at the G2/M border and released for 40min, 120min and 240min.

**Supplementary Table 6. TF footprints analysis of distal CREs gaining chromatin accessibility upon acute SOX9 depletion; related to Supplementary Figure 4**

The table reports the results from TOBIAS analysis at distal CREs gaining chromatin accessibility at the G2/M border and released for 40min, 120min and 240min.

**Supplementary Table 7. Functional terms associated with depletion of SOX9 prior to mitosis; related to Figure 6**

The table reports the gene sets used for Gene Set Enrichment analysis (GSEA) at t=240min after release from G2/M arrest and with or without prior depletion of SOX9.
